# Molecular architecture of black widow spider neurotoxins

**DOI:** 10.1101/2021.04.19.440504

**Authors:** Minghao Chen, Daniel Blum, Lena Engelhard, Stefan Raunser, Richard Wagner, Christos Gatsogiannis

## Abstract

Latrotoxins (LaTXs) are presynaptic pore-forming neurotoxins found in the venom of *Latrodectus* spiders. The venom contains a toxic cocktail of seven LaTXs, with one of them targeting vertebrates (α-latrotoxin (α-LTX)), five specialized on insects (α, β, γ, δ, ɛ-latroinsectotoxins (LITs), and one on crustaceans (α-latrocrustatoxin (α-LCT)). LaTXs bind to specific receptors on the surface of neuronal cells, inducing the release of neurotransmitters either by directly stimulating exocytosis or by forming Ca^2+^-conductive tetrameric pores in the membrane. Despite extensive studies in the past decades, a high-resolution structure of a LaTX is not yet available and the precise mechanism of LaTX action remains unclear.

Here, we report cryoEM structures of the α-LCT monomer and the δ-LIT dimer. The structures reveal that LaTXs are organized in four domains. A C-terminal domain of ankyrin-like repeats shields a central membrane insertion domain of six parallel α-helices. Both domains are flexibly linked via an N-terminal α-helical domain and a small β-sheet domain. A comparison between the structures suggests that oligomerization involves major conformational changes in LaTXs with longer C-terminal domains. Based on our data we propose a cyclic mechanism of oligomerization, taking place prior membrane insertion. Both recombinant α-LCT and δ-LIT form channels in artificial membrane bilayers, that are stabilized by Ca^2+^ ions and allow calcium flux at negative membrane potentials. Our comparative analysis between α-LCT and δ-LIT provides first crucial insights towards understanding the molecular mechanism of the LaTX family.

## Introduction

Latrotoxins (LaTXs) are potent high molecular weight neurotoxins from the venom of black widow spiders. The venom contains an arsenal of phylum-specific toxins, including one vertebrate-specific toxin, α-latrotoxin (α-LTX)^1^, five highly specific insecticidal toxins (α-, β-, γ-, δ-, and ɛ-latroinsectotoxin (LITs))^2,3^, and one crustacean-specific toxin, α-latrocrustatoxin (α-LCT)^3,4^. The vertebrate specific α-LTX causes a clinical syndrome named lactrodectism upon a venomous bite to humans, which is fortunately rarely life-threatening but often characterized by severe muscle cramps and numerous other side effects such as hypertension, sweating, and vomiting^5,6^.

LaTXs are produced as ~160 kDa inactive precursor polypeptides in venom glands and secreted into the gland lumen. There the final mature 130 kDa toxin is produced by proteolytic processing at two furin sites and cleavage of a N-terminal signal peptide and a C-terminal inhibitory domain^7,8^. Most of the physiological and molecular biological researches to date have been carried out using the vertebrate specific toxin α-LTX.

α-LTXs have been shown to form cation-selective pores upon binding to specific receptors on the presynaptic membrane and induce Ca^2+^influx, mimicking thereby physiological voltage-dependent calcium channels^9,10^. Ca^2+^influx activates the exocytosis machinery^11^and triggers massive release of neurotransmitters. α-LTX was shown to form also pores on artificial lipid bilayers, which have high conductance for monovalent and divalent cations such as K^+^, Na^+^, Ca^2+^, and Mg^2+^, but are blocked by transition metals and trivalent ions such as Cd^2+^and La^3+^ ^12–16^. Efficient incorporation into biological membranes, strictly relies however on the presence of specific receptors^17–19^.

To date, three receptors for α-LTX have been isolated, *i.e*., the cell adhesion protein neurexin (NRX) which binds to the latrotoxin in a Ca^2+^-dependent manner^20–22^, the G protein-coupled receptor latrophilin (LPHN or CIRL, stands for Calcium-Independent Receptor of Latrotoxin)^23,24^and the receptor-like protein tyrosine phosphatase σ (PTPσ)^25^. NRX and PTPσ are suggested to provide with regard to α-LTX only a platform for binding and subsequent pore formation events^25–29^. Intriguingly, Ca^2+^-independent binding to receptor LPHN does not involve in contrast oligomerization and channel formation, but direct downstream stimulation of the synaptic fusion machinery^27,30–32^.

The channel dependent and independent functions of α-LTX have attracted the attention of neurobiologists for several decades, studying the effects of α-LTX on neurotransmitter release and mechanisms underlying synaptic plasticity. The α-LTX variant LTX^N4C^ ^33^, which lacks the ability of pore-formation but retains the full binding affinity to receptors, played a key role in the investigations of α-LTX action. Today, α-LTX is an indispensable tool for stimulating exocytosis of nerve and endocrine cells^29,34–36^. α-LTXs are furthermore considered to antagonize botulinum poisoning and attenuate the neuromuscular paralysis via synapse remodeling^37^. The surprising structural homology of α-LTX to the glucogen-like peptide-1 (GLP1) receptor might also open opportunities for pharmacological applications in blood glucose normalization and reversion of neuropathies^38^.

Invertebrate LaTXs are less well understood, but considered as promising candidates for the development of novel bio-pesticides. Orthologues of the three receptor classes shown to bind α-LTX are also present in insects^39^. To date, four LaTXs have been cloned, including α-LTX^7^, α-LIT^40^, δ-LIT^41^, and α-LCT^42^. Despite their high specificity, the different LaTXs display a 30-60% sequence identity and are expected to share an overall similar domain organization and membrane insertion mechanism. Low resolution 3D maps (14-18 Å) of the α-LTX dimer, α-LTX tetramer, and δ-LIT monomer were previously determined using single particle negative stain and cryoEM^39,43^, suggesting indeed an overall similar architecture of the different members of the LaTX family.

A structural and mechanistic understanding of LaTX function is a significant priority for the development of novel anti-toxin therapeutics and/or insecticides. However, a high-resolution structure of a LaTX, which is a prerequisite for the understanding of LaTXs’ mechanism of action at molecular detail, has been missing. Here we present a 4.0 Å cryoEM structure of the α-LCT monomer and a 4.6 Å cryoEM structure of the δ-LIT dimer revealing the molecular architecture of LaTX neurotoxins as well as the molecular details of their oligomerization mechanism prior to membrane insertion. In addition we characterized the principal basic pore characteristics of α-LCT, the precursor δ-LIT and δ-LIT channels after reconstitution into planar lipid bilayer.

## Results

### CryoEM structure determination of α-LCT

We recombinantly expressed the mature α-LCT (amino acids 16-1240) (**Fig. 1a and Supplementary Fig. 1**) in insect cells using the MultiBac system and purified it using a combination of affinity and size exclusion chromatography to obtain a monodisperse sample for cryoEM single particle analysis (**Supplementary Fig. 2a, b**). The cryoEM sample showed a homogeneous set of characteristic G-shaped flat particles, corresponding to soluble monomers of the 130 kDa mature α-LCT complex (prepore state; before membrane insertion). The G-shaped particle is composed of a C-like curved region, corresponding to the long C-terminal domain of ankyrin-like repeats (ARs), that is engulfing a central compact “head” region (**Fig. 1b, Supplementary Fig. 2c**).

**Figure 1.**
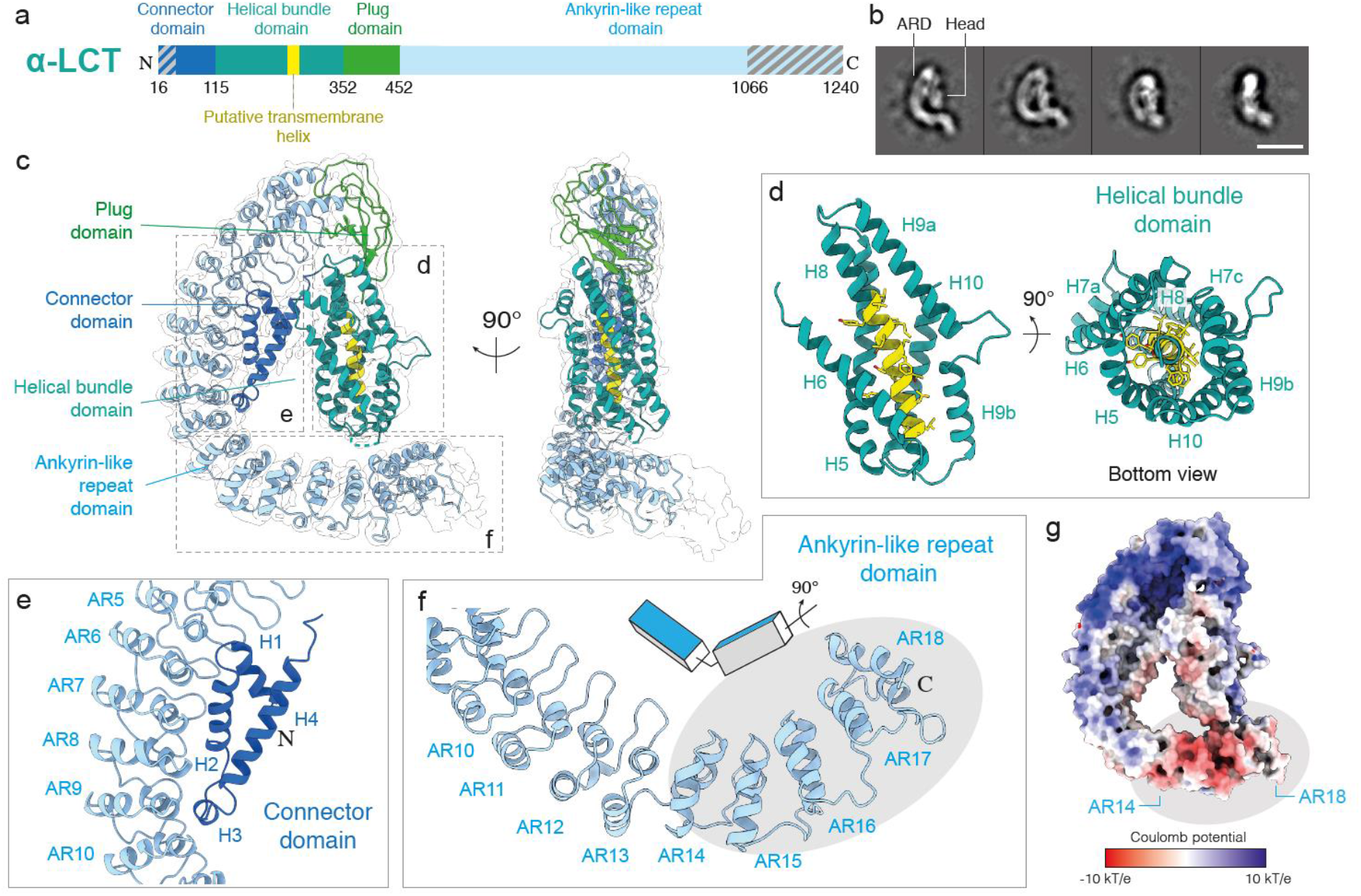
Structure of α-LCT monomer. **a**, Domain organization of mature α-LCT. Gray diagonal lines indicate regions not resolved in the cryoEM density. **b**, Representative reference free two-dimensional class averages. Scale bar: 10 nm. **c**, Side views of the α-LCT monomer superposed with the EM map (transparent). Domains are depicted in the same colors as in a. **d**, Close-up view of the helical bundle domain. The front helix (H7) is not shown in the left image for clarity. **e**, Close-up view of the interface between CD (H1-3) and ARD (AR5-10). **f**, Close-up view of the ARD C-terminal tail. The gray ellipse indicates the last five ARs (AR14-18). Note the change in orientation **g**, Electrostatic potential calculated in APBS. Red: −10 kT/e; Blue: +10 kT/e. PD:plug domain; HBD:helical bundle domain; CD: connector domain; ARD: ankyrin-like repeat domain

Subsequent image processing and 3D classifications revealed an inherent flexibility between both regions. The N-terminal head is orientated perpendicular and in close vicinity to the tail of the AR region in the best resolved class (“compact” conformation), but shows a continuous movement and is tilted away the tail of the AR-domain in less well resolved classes, resulting in less compact conformations (**Supplementary Fig. 2d,e, 3a**).

The α-LCT monomer in the “compact” conformation adopts a flat architecture and is 130 Å long and 30 Å wide. The path of the polypeptide is clear in the density map, allowing us to build an atomic model covering 85% of the sequence of the molecule (residues 48-1066, except two disordered loop regions 226-232, 349-360) (**Supplementary Fig. 4a**). We deleted side-chain atoms beyond Cβ in regions where side-chain density was only rarely evident. The C-terminal end of the ARs domain including the last four ankyrin repeats is not resolved. The local resolution is highest in the N-terminal head region (**Supplementary Fig. 2h**).

### Architecture of the soluble α-LCT monomer

The resulting atomic model reveals that the N-terminal head region is composed of three domains: a four-helix domain at the N-terminal end, termed here as “connector” domain (residues 48-115); a central helical bundle domain (residues 116-352), and a short β-sheet domain, linking the helical bundle domain with the AR domain, termed here as “plug” domain (residues 353-452) (**Fig. 1a,c, Supplementary Fig. 5, Movie 1**).

The helical bundle domain shows a novel fold of a six-helix bundle, with five parallel aligned helices assembling into a cylindrical structure (H5-7, H9-10), encircling a central α-helix (H8) (**Fig. 1d**). The conserved helix H8 contains many hydrophobic residues and is predicted to act as transmembrane region of the tetrameric LaTX pore (**Supplementary Fig. 6d**). The surrounding helices shield the hydrophobic surface of H8 within the cylindrical bundle and protect it from the aqueous environment (**Supplementary Fig. 7a,b**). The helical bundle domain is expected to undergo severe conformational changes during pore formation to allow exposure of the transmembrane helix H8 and transition of the toxin from a soluble monomer to a transmembrane tetramer. Interestingly, helices H7 and H9 are kinked and interrupted by two (residues 191-196, 203-211) and one (residues 296-304) short loops, respectively. Such short breaks interrupting long α-helices in close vicinity to the putative transmembrane regions, were shown in other α-helical pore forming toxins to provide the necessary flexibility for major conformational changes towards membrane insertion^44^.

The long and curved C-shaped C-terminal AR domain consists of 22 ankyrin-like repeats, accounting for two thirds of the sequence of α-LCT. In total, the first 18 out of 22 ankyrin-like repeats (ARs) were resolved in the present map. Interestingly, there is a redirection of orientation of the ARs at the loop (residues 890-902) connecting AR13 and AR14. ARs14-18 are rotated almost 90 degrees compared to ARs1-13 along the long axis of the domain (**Fig. 1f**). This arrangement goes along with a characteristic bipolar charge distribution, with ARs1-13 dominantly positively charged and the tail of ARs14-18 displaying a prominent negatively charged patch (**Fig. 1g, Supplementary Fig. 6a**).

The “connector” domain at the N-terminal end forms the “lower” interface connecting the central helical bundle with the AR-domain (**Fig. 1c,e**). Helices H1, H2, H4 and the short helix H3 of the “connector “domain, assemble into a flat triangle structure, which is attached to the inner curved surface of the AR-domain and interacts with ARs 6-10. Most interaction surface to AR-domain is provided by H2, whereas the short helix H3 is directly positioned in close proximity to the loop connecting AR9 and AR10 and parallel aligned to the helices of AR9 and AR10 (**Fig. 1e**). The residues involved in this interface are only conserved in the AR-, not in the “connector” domain (**Supplementary Fig. 7c,d**). They are mainly hydrophobic, suggesting a major role of hydrophobic interactions. Helix H2 of the “connector” domain is hydrophobic in our structure and interestingly, this helix has been also predicted as the second transmembrane region of the insecticidal δ-LIT (**Supplementary Fig. 6h**), but not predicted as such from the sequences of other latrotoxin family members, such as α-LCT (**Supplementary Fig. 6d**), or α-LTX (data not shown).

The “plug”-domain covalently links the primary sequence of the helical bundle with the AR-domain, positioning the helical bundle directly below AR1 (“upper interface”) (**Fig. 1c, Supplementary Fig. 7e-g**). The “plug”-domain is organized in two layers: a region of several flexible loops and a core region of four β-strands that is attached to H5 and a short loop between H8 and H9 of the helical bundle (**Supplementary Fig. 7h**). The “plug”-domain plays an important role in the oligomerization of the complex prior complex formation, which will be discussed in detail in the next section.

### CryoEM structure of inactive precursor soluble δ-LIT dimer

In subsequent experiments, we were not able to induce oligomerization of α-LCT and trigger insertion into liposomes for further visualization of pore formation events, as previously described for α-LTX^43,45^. To provide further insights into the LaTX family, we then focused on the insecticidal δ-LIT. As expected, mature δ-LIT was toxic for our insect cell cultures and therefore, we expressed, purified, and subjected to cryoEM analysis the precursor uncleaved inactive toxin. The precursor toxin contains an additional signal peptide (residues 1-28) and an inhibitory domain (α-LCT residues 1037-1214), compared to the matured form (**Fig. 2a**). Albeit we performed the cryoEM analysis with the same procedure used for mature α-LCT, subsequent processing did not only reveal G-shaped monomers, as was the case for mature α-LCT, but also higher order oligomers. In particular, reference free 2D classifications revealed 40% monomers, 50% dimers, 5% trimers and 5% tetramers (**Fig. 2b, Supplementary Fig 8a-c**). Interestingly, such particle populations of oligomers were not observed in negative-stain EM (**Supplementary Fig. 9a,b**), indicating that the interactions are rather weak and dilution of the sample which is necessary for negative-stain EM, as well as the low pH of the stain might induce dissociation of the oligomers. We were finally able to obtain a map of the δ-LIT dimer at 4.6 Å average resolution from 81,192 particles, with no symmetry imposed (**Supplementary Fig. 8d-f**).

**Figure 2.**
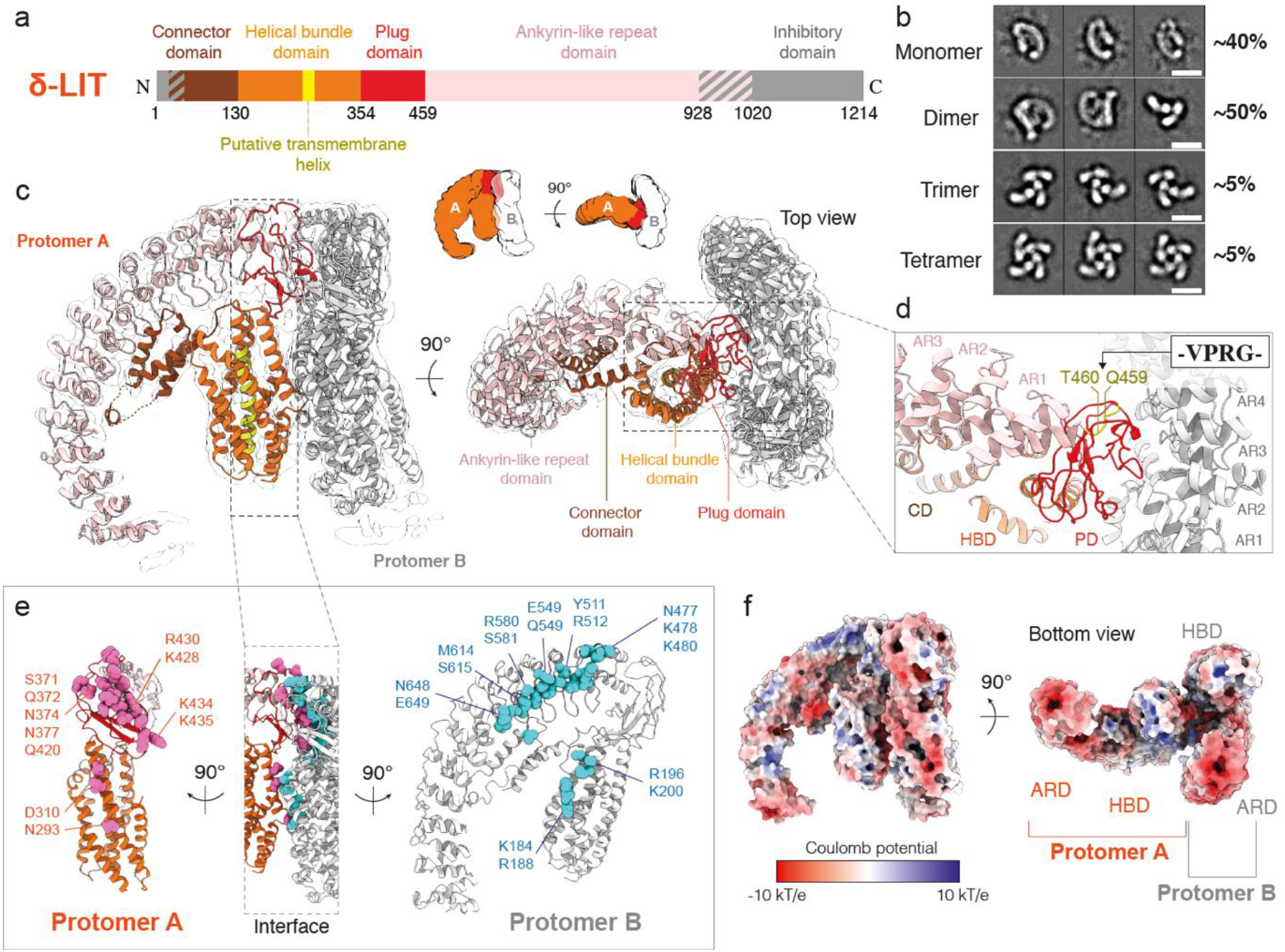
Structure of δ-LIT dimer. **a**, Domain organization of full-length precursor δ-LIT. Disordered regions in the cryoEM map are indicated in gray diagonal lines and boxes. **b**, Representative two-dimensional class averages of each oligomeric state. Scale bar: 100 Å. **c**, Side and top view of the δ-LIT dimer superposed with the EM map (transparent). Domains of protomer A are depicted in the same colors as in a; protomer B is colored in gray. **d**, Close-up view of the PD-ARD dimerization interface. The position of the four amino acid insertion variant (VPRG) is indicated and colored in yellow. **e**, Side view of the dimerization interface. Protomers A and B are rotated 90° to left and right, for better clarity. Polar and charged candidate residues (<5 Å to the opposite protomer) are shown as spheres and colored in cyan and pink. **f**, Electrostatic potential calculated in APBS. Red: −10 kT/e; Blue: +10 kT/e. PD:plug doman; HBD:helical bundle domain; CD: connector domain; ARD: ankyrin-like repeat domain

A 3D reconstruction of the δ-LIT monomer was unfortunately not possible, due to preferred orientation of the single particles. Based on the structure of α-LCT monomer, we were able to build a molecular model of the δ-LIT dimer, including residues 50-928 for both protomers (**Fig. 2c, Supplementary Fig. 4b, Movie 2**). As expected, the structure of the protomer of δ-LIT is very similar to that of α-LCT (sequence identity 39%), showing the characteristic G-shaped architecture and domain organization. δ-LIT displays however a shorter repetitive C-terminal AR-domain, containing only 15 ARs instead of 22 in α-LCT.

**Figure 3.**
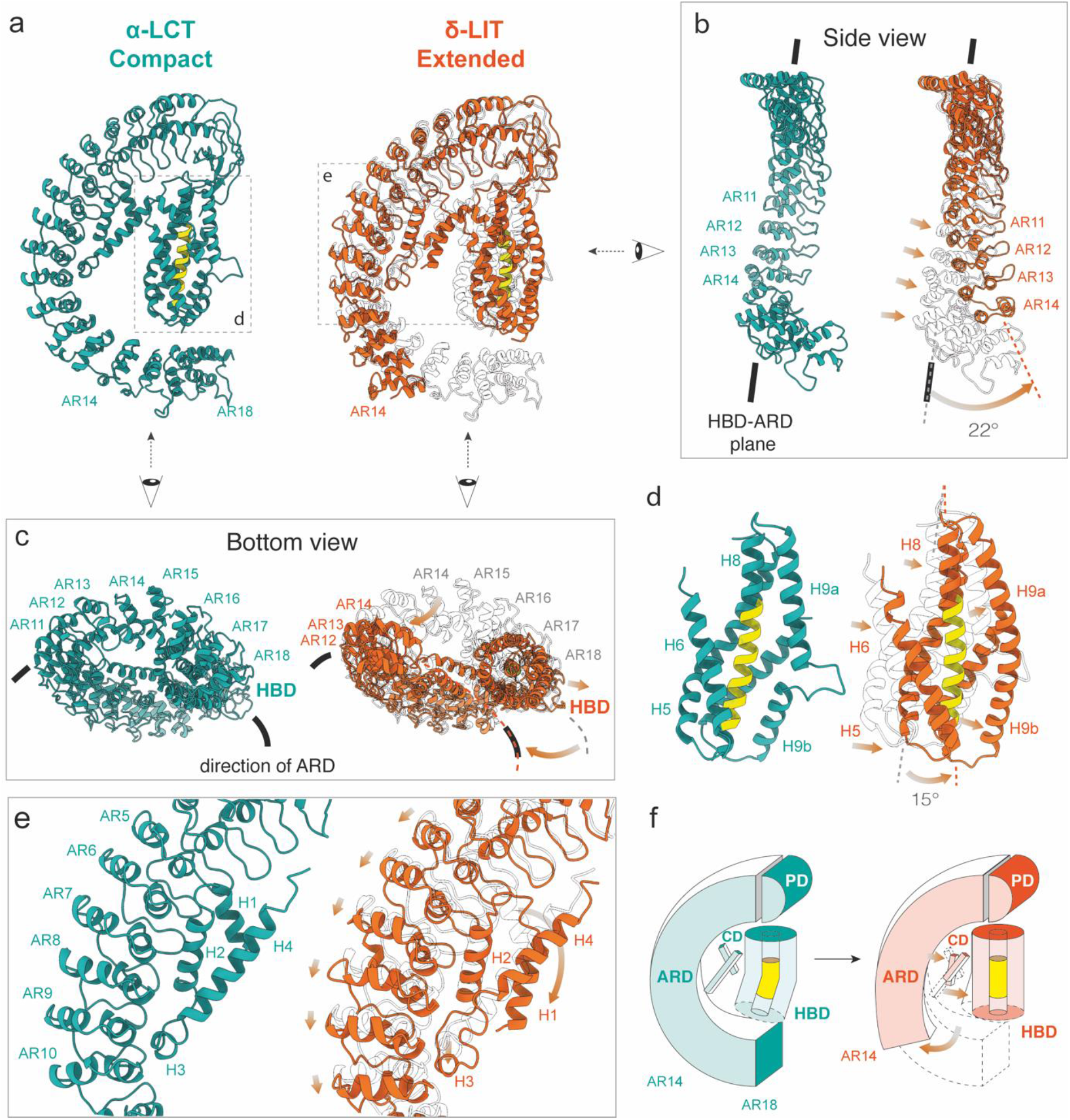
Conformational changes during dimerization. **a** Side-by-side comparison of α-LCT (compact state, sea green) and d-LIT (extended state, extracted from the dimer, orange) superposed with α-LCT (transparent). **b**, Front view of the ARD. Arrows indicate domain motion during dimerization **c**, Bottom views of the compact and extended states. The bottom part of the cylindrical (HBD) is exposed in the extended state. **d**, Magnified view of the HBDs. The front helix (H7) is not shown for clarity. The H9a-H9b loop folds helically to complete H9 in the extended state. **e**, Magnified view of the interface between the connector- and the AR-domain **f**, the schematic diagram illustrates the conformational change. PD:plug domain; HBD:helical bundle domain; CD: connector domain; ARD: ankyrin-like repeat domain

**Figure 4.**
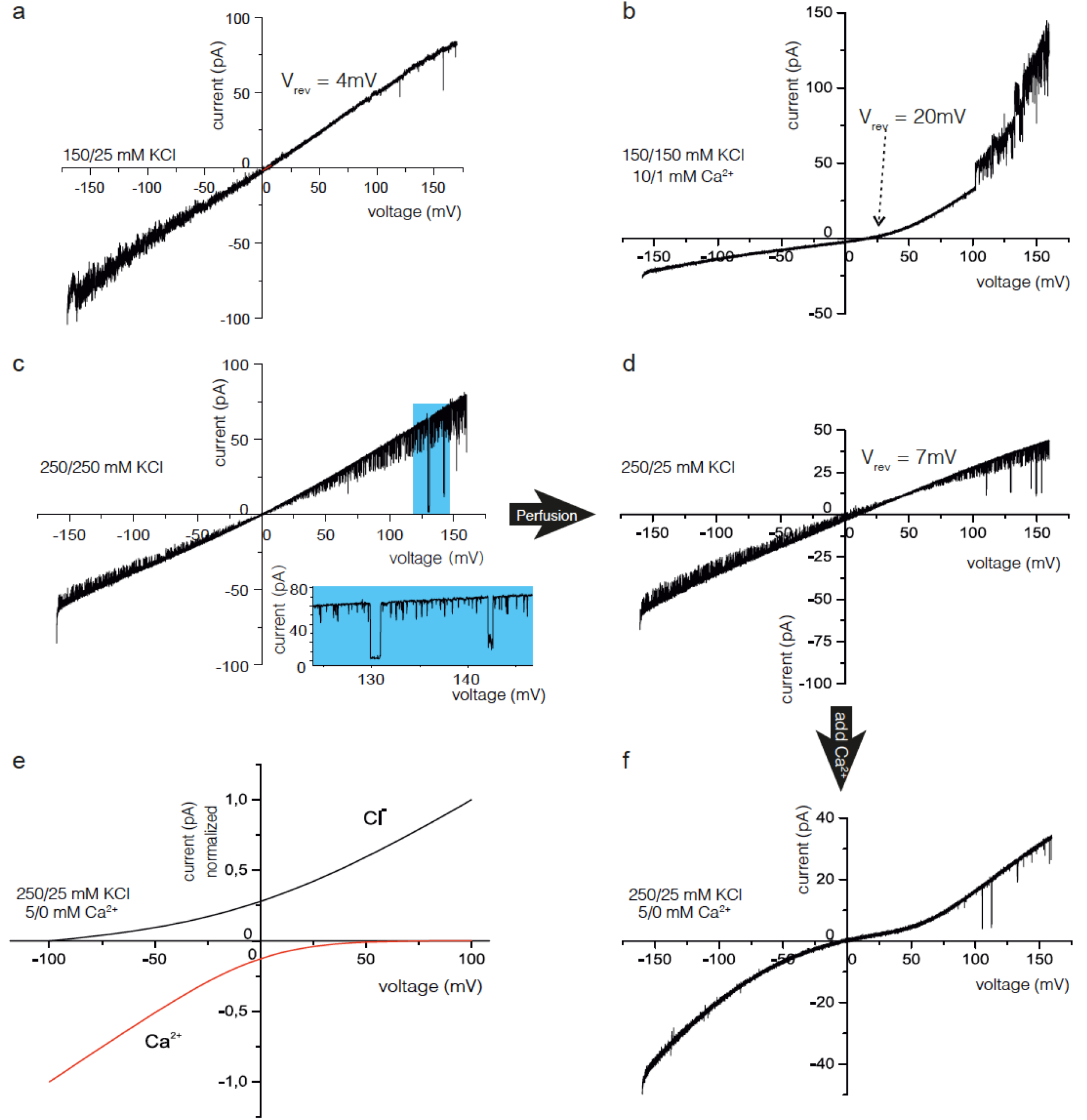
Calcium induced ion channel properties of reconstituted δ-LIT variants within lipid bilayers. Current-Voltage ramp recordings at different cis/trans buffer conditions. **a**, Precursor δ-LIT shows a noisy linear current-voltage relation with reversal potential of V_rev_ = +4mV, indicating a slight cation selective channel (*P_K^+^_/P_Cl^−^_* =1.25). **b**, With symmetric KCl and added asymmetric calcium the precursor δ-LIT channel displays a V_rev_ = +20 mV, presenting now a highly Ca^2+^ selective channel (*P_Ca^2+^_/P_Cl^−^_* =18), demonstrating a dramatic property change induced by the presence of *Ca*^2+^-ions. **c**, The current-voltage relation of the mature δ-LIT is again linear but with low current noise and frequent, defined gating from the open towards the closed state. (displayed in extension box). **d**, Next trans chamber perfusion within the same experiment. The V_rev_ = 7 mV value discloses cation selective properties (*P_K^+^_/P_Cl^−^_*=1.47) preserved within the mature variant. **e**, With asymmetric KCl and cis added calcium (no calcium addition trans) the precursor δ-LIT channel displays an asymptotic sine current-voltage relation with a zero-current crossing at V_rev_ = 0 mV. Extrapolation of the tangent to zero net current yields *V_rev_* =≈ 40 *mV* and a very high calcium selectivity of the mature δ-LIT channel (*P_Ca^2+^_P_K^+^_/P_Cl^−^_* ≌ 600/1.47/1). **f**, Calculated GHK current-voltage relation using the experimental relative permeabilities from (e) and the experimental concentration of *Ca*^2+^, *K*^+^ and *Cl*^−^ions in the cis and trans compartment (see Supplemental Information for Details).

The path of the polypeptide is clear for all four domains, i.e., connector domain, helical bundle domain, plug domain, and AR-domain, except for a few loop regions (residues 91-92, 99-105, 237-245, 355-365) in protomer A and (237-244, 356-361) in protomer B, the last (15^th^) AR and the complete signal peptide and inhibitory domain in both protomers (**Fig. 2c**). These structural regions were not resolved in the cryoEM density map, probably because they are disordered or highly flexible. Due to limited resolution, we deleted most of the side-chain atoms beyond Cβ in the molecular model unless the electron density was sufficiently clear with regard to bulky side chains.

The δ-LIT dimer is formed with the two protomers rotated 90 degrees relative to each other and the “plug” domain of protomer A plugging from the side into a cleft formed by ARs 1-6 of protomer B (**Fig. 2c**). The protomers A and B are in basically the same conformation, but the small differences, mainly due to the flexibility of the helical bundle domain, precluded successful C2 symmetrization of the particle (**Supplementary Fig. 3e**).

The “plug”-domain has a hemispherical architecture matching the cleft formed by the 1^st^-6^th^ARs of the AR-domain (**Fig. 2d**), suggesting an induced fit and shape complementarity as the basis for the interaction. We found clusters of 17 polar and charged residues on the “plug”-domain of protomer A and 18 polar and charged residues on the cleft of protomer B, that may be involved in this interface (**Figure 2e, Supplementary Table 1**). In previous studies, the α-LTX variant LTX^N4C^with an insertion of four amino acids (VPRG) at the linker peptide connecting the plug domain with the AR-domain^33^, was shown to retain its binding affinity to receptors, but lose its ability to oligomerize into tetramers and form pores^27^. This variant played a key role in understanding of the dual mode of action of LaTXs. The corresponding position of this insertion is highlighted on the molecular model of δ-LIT (**Fig. 2d**). This insertion is not positioned directly at the dimerization interface, but in close proximity to the loops of “plug”-domain involved in the interaction and might thus disturb the overall shape complementarity and/or induce a shift of positions and a mismatch between the residues involved in this interface, blocking thereby dimerization.

Besides the main interaction between the “plug”-domain of protomer A and ARs 1-6 of protomer B, there is a less pronounced interaction between the helical bundle domains of both protomers, involving H9 of protomer A and H6/H7a of protomer B (**Fig. 2e, Supplementary Table 1**). Whereas H9 is interrupted in protomer A and in the structure of α-LCT by a short loop in its middle, in protomer B (thus most probably upon dimer formation), this loop folds helically to straighten and complete H9 (**Supplementary Fig. 3f, Supplementary Fig. 4b**). Interestingly, because δ-LIT is lacking seven terminal ARs, the surface of the AR-domain does not display a clear bipolar charge distribution as in α-LCT (**Fig. 2f, Supplementary Fig. 6e**). However, in both LaTXs, the terminal tail of the AR-domain is clearly negatively charged (**Supplementary Fig. 6a, e**).

### Structural differences between δ-LIT and α-LCT

In comparison to the “compact” conformation of the α-LCT monomer, both protomers of the δ-LIT dimer show a different, rather “extended” conformation: the outermost helix H9 and the overall helical bundle domain are straightened, the helical bundle of each protomer is further tilted 15 degrees away the long axis of the AR-domain and the distance between the helical bundle domain and the shorter AR-domain is substantially longer. Notably, the AR-domain of δ-LIT is significantly less curved (**Fig. 3a**) and its C-terminal tail is further’twisted’ and positioned outside the helical bundle domain and ankyrin-like repeat domain (HBD-ARD) plane (**Fig. 3b**). As a consequence, the bottom part of the helical bundle domain becomes in this case exposed (**Fig. 3c**).

The enlarged distance between AR-domain and helical bundle domain, requires also a more extended conformation of the connector domain, which is further stretched in δ-LIT, but still bridges both domains. This results into repositioning of H1 and unfolding of the lower half of H4 of the connector domain (**Fig. 3e**). These differences between the compact α-LCT and the extended δ-LIT are summarized in Movie 3 and a schematic diagram in **Fig. 3f**. Interestingly, the less well resolved 3D class of α-LCT can be considered as flexible intermediate between both structures (**Supplementary Fig 3g, h, Movie3**).

We were not able to obtain a structure of the δ-LIT monomer, but the respective 2D class averages, suggest also flexibility for the helical bundle domain and possibly a mixture of different compact, intermediate and extended conformations (**Fig. 2b**). α-LCT appears in general however more compact, due to the seven additional terminal ARs, allowing additional interactions between the AR-domain and the flexible central helical bundle. This might explain the different oligomerization properties observed for both molecules under identical cryoEM conditions. The directional change observed however in the AR-domain of the “extended” δ-LIT protomer, would also result in exposure of the helical bundle domains, even for LaTXs with significantly longer AR-domains such as α-LCT or α-LTX (**Fig. 3c**, right panel).

It should be noticed that the previous low resolution EM studies on α-LTX and δ-LIT show an overall different LaTX architecture. The previous 2D cryoEM class averages of α-LTX^43^and the present 2D classes of δ-LIT dimers and tetramers (**Fig. 2b**) display however clear similarities, indicating structure conservation. Taking in addition in account the high sequence similarity within the LaTX family, we rather conclude that the earlier LaTX reconstructions determined more than two decades ago, are apparently affected by the previous bottlenecks of the technique and the significantly lower signal to noise ratio in the cryoEM micrographs. The members of the LaTX family share an overall common domain organization and architecture.

### Electrophysiological characteristics of the pore forming precursor δ-LIT in comparison with the mature δ-LIT and α-LCT

We further performed electrophysiology studies to demonstrate pore-formation activity of the recombinant proteins and provide a detailed characterization for the less well studied invertebrate LaTX channels. One important aspect herein is the role of the C-terminal domain in channel formation, since its cleavage is required for activation of the toxins. Therefore, we additionally prepared precursor full length δ-LIT for further functional comparisons with mature truncated δ-LIT and α-LCT. The inherent cleavage sites were however not recognized by furin protease (data not shown) and therefore we inserted two additional cleavage sites into the sequence of precursor δ-LIT, before the C-terminal-(residue 29) and after the AR-domain

(residue 1019) followed by proteolysis treatment after expression (**Supplementary Fig. 9c,d**). The two mature toxins and precursor δ-LIT were then reconstituted into planar lipid bilayer and voltage depended membrane currents (V_mem_, membrane potential) were recorded at single channel resolution as previously described ^46^.

Precursor δ-LIT spontaneously inserted into the lipid bilayer and formed open channels as obvious from the observed large voltage induced currents (**Supplementary Fig. 10a,c**). With asymmetric (cis/trans) 150/25 mM KCl buffer conditions, the reversal potential (V_rev_) was V_rev_ = 4 ± 1 mV yielding P_K+_ / P_cl^−^_ =1.25^47,48^, demonstrating only marginal selectivity to K^+^ over Cl^−^ ions (**Fig. 4a**). However, upon addition of 10/1 mM CaCl_2_ (cis/trans)) gradient in symmetrical 150/150 mM KCl buffer, we observed an approximate five-fold increase in the reversal potential (V_rev_ = 20 ± 2,5 *m*V (**Fig. 4b**)) revealing that the precursor δ-LIT voltage activated channel preferentially conducts *Ca*^2+^ ions (P_ca^2+^_ /P_cl^−^_=18) and the current-voltage relation became rectifying. Moreover, calcium stabilized the channel, visible by a significant reduction in the current noise. The mean main conductance as determined from all point current histograms at different voltages was 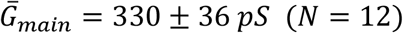

The current voltage relations of membrane inserted mature δ-LIT in symmetrical 250/250 mM KCl buffer showed a linear course with high gating activity(**Fig. 4c**). In contrast to the precursor variant, we observed low noise currents with structured gating which could be resolved in amplitude and time. Similarly to the precursor-, mature δ-LIT channels display also only a slight cation selectivity (P_K^+^_ /P_cl^−^_ =1.47, **Fig. 4d**) after establishing the 250/25mM KCl gradient. Addition of 5 mM Ca^2+^ ions to the cis side of the bilayer changed however the properties of the mature δ-LIT channel completely. The asymptotic-sine course of the current-voltage relation increased at both negative and positive command voltages (V_*cmd*_) while crossing the zero voltage with zero net current (**Fig. 4f**). To further analyze this, we applied the widely used Goldman-Hodgkin-Katz (GHK) approach^47,48 49^(see Supplemental for details). It turned out that the course of the current-voltage relation in **Figure 4f** can be explained if the currents at negative V_*cmd*_ are carried from cis to trans mainly by Ca^2+^-ions and at positive V_*cmd*_ predominantly by Cl^−^ions (**Fig. 4e**; see Supplemental for details). The analysis of current traces revealed two open channel states with mean amplitudes of 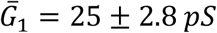 and 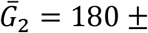 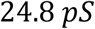 corresponding to a pore restriction diameter of about 0.8nm^48,50^(**Supplementary Fig. 10e,f**) (N=15) significantly smaller than the one calculated for the precursor δ-LIT channel (~1.5nm see **Supplementary Figure 10c**). Thus the mature δ-LIT channel appears to form a denser, stabilized conformation compared to its precursor variant. Additionally, the mature δ-LIT seems to act like a complete rectifier, which in the presence of Ca^2+^, depending on membrane polarization, allows mainly either flux of calcium (−V_*mem*_) or chloride currents (+V_*mem*_) from cis to trans. In this context, it seems important that both the precursor δ-LIT and the mature δ-LIT are incorporated into the bilayer predominantly in a unidirectional manner, with a regulative Ca^2+^-binding site on the cis side. Strong rectifying current-voltage relations were observed in all bilayer experiments in the presence of Ca^2+^-ions added on the cis side (**Figure 4b, f**).

Surprisingly, mature α-LCT incorporates into the bilayer membrane and forms channels only in the presence of Ca^2+^-ions (cis) (**Supplementary Fig 10g**). The analysis of the single channel traces (**Supplementary Fig 10h,i**) revealed, similar to mature δ-LIT, two distinct open channel states 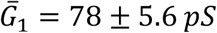 and 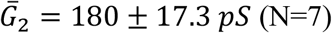 (**Supplementary Fig 10i**). Beside the 10/1mM CaCl_2_ gradient, the α-LCT channel does not show, in contrast to δ-LIT, any preference for the involved anions and cations (V_*rev*_ = 0 ± 2.1 mV (**Supplementary Fig. 10g**). In comparison to mature δ-LIT (**Fig. 4f**), the α-LCT channel displays a higher current noise level in the recordings and surprisingly, considering the slightly asymptotic sine course of the current-voltage relation (**Supplementary Fig 10g**), the α-LCT apparently also harbors rectifying properties, making it likely that rectified currents of Ca^2+^-ion and Cl^−^ —ions may convey similar to mature δ-LIT.

To sum up, Ca^2+^-ions appear to further stabilize precursor and mature LaTX oligomers after incorporation into the membrane resulting in a rectifying calcium selective channel allowing calcium flux at negative membrane potentials. High Ca^2+^permeability was previously also shown for native δ-LIT channels in locust muscle membrane and artificial bilayer, but not for truncated recombinant variants^41^. Provided that as a result of the predominantly unidirectional insertion the cis side corresponds to the cytosolic side of the respective channel, δ-LIT and α- LCT display ion channel characteristics similar to Ca^2+^-release channels^51^.

### Formation of an insertion-competent prepore complex

The reference free 2D class averages of monomers, dimers, trimers and tetramers identified in the δ-LIT cryoEM dataset (**Fig. 2b**) suggest a general sequential oligomerization mechanism of LaTXs towards the formation of a soluble tetrameric complex prior membrane insertion. The bent cleft at the upper part of the AR-domain of a protomer is employed as a binding site for the hemispherical plug domain of an adjacent protomer, with both molecules rotated 90 degrees relative to each other. Four LaTX molecules dock into each other in a sequential circular ring-fashion form via 1/2/3-mers to eventually form tetramers (**Supplementary Fig. 11a**). This is to our knowledge the first description of a LaTX intermediate trimer, suggesting that the tetramer is not exclusively formed upon assembly of two dimers, as previously proposed^45^.

Based on our data, we generated a 3D model of the tetramer by arranging the δ-LIT dimer (“extended state”) according the 2D class averages of the top view of the tetramer. The resulting 3D model of the tetramer is approximately 140×140 Å large and has a height of approximately 120 Å, with an overall striking configuration resembling a “four-finger crane claw”, with each curved AR-domain resembling a finger of the crane claw (**Fig. 5a**).

In the resulting model, the four cylindrical helical bundle domains are surrounding a central 10 Å diameter channel, which agrees excellent with the 2D class averages of the soluble δ-LIT tetramer (**Fig. 2b**) and the previous 2D class averages of the soluble α-LTX tetramer^43^. The putative transmembrane helices are still however completely shielded from the aqueous environment within the respective helical bundle domains. Nevertheless, the bottom part of the helical bundle domains is exposed in this arrangement. We therefore propose that the tetramer, shown here, composed of “extended” protomers, resembles an insertion-competent prepore state of the toxin.

Assembling the tetramer in a similar manner from α-LCT “compact” protomers with 22 ARs results instead into a completely closed “crane claw” (**Supplementary Fig. 11b**). In this scenario, the AR-domains shield the central helical bundle domains from both sides and the central channel is closed. The resulting compact structure does not match however the 2D class averages of LaTX tetramers (**Fig. 2b** and Orlova *et. al.*^43^) and there are severe clashes between the AR-domains of adjacent subunits. Note that in previous 2D class averages of α-LTX tetramers with 22 ARs^43^, the central helical bundle domains are also exposed, suggesting that the single protomers are in the “extended” conformation. This suggests that α-LCT has to undergo a conformational switch from the compact to extended conformation during the oligomerization process and not after formation of the tetramer. Low resolution 3D classes of the α-LCT monomer resembling intermediate states between the compact and the extended conformation further support this scenario (**Supplementary Fig. 3g,h**, **Supplementary Movie 3**). α-LCT shows an intriguing bipolar charge contribution, which is apparently present also in α-LTX which has also 22 ARs, but however less pronounced in δ-LIT with 15 ARs (**Supplementary Fig. 12, Supplementary Fig. 6a,e**). The “claw tips” (lower ends of the AR-domains) are however clearly negatively charged in all LaTXs.

## Discussion

### Oligomerization characteristics and function of the inhibitory domain

Albeit extensive efforts, also in presence of artificial membranes, we were not able to determine *in vitro*, in absence of receptors, factors controlling the oligomerization and subsequent pore formation process efficiently for further visualization of the pore formation events. Interestingly, we observed the various populations of prepore oligomers of precursor δ-LIT (5% of the particles) only in cryoEM samples (**Supplementary Fig. 8b,c**) but not during negative stain EM analysis (**Supplementary Fig. 9a,b**) or other complementary methods (e.g. SEC-MALS, data not shown). The oligomerization occurred in buffer containing EDTA, and the substitution of EDTA with Mg^2+^or Ca^2+^did not affect the sample (data not shown). This observation corresponds to a previously reported study^45^, suggesting oligomerization of δ-LIT as a process independent of divalent cations. Considering in addition that active α-LCT formed exclusively stable monomers throughout our experiments, as well as previous studies suggesting that α-LTX exists mainly in its dimerized or tetramerized form, we conclude that different LaTXs display different oligomerization characteristics. Indeed, although the dimerization interface suggests a general induced fit mechanism in LaTXs, the surface of the involved AR-domain cleft is not highly conserved in the LaTX family (**Supplementary Fig. 1** and **Supplementary Fig.6c, g**).

The electrophysiological analysis at single channel resolution allowed us to confirm the tetramerization process for both recombinant expressed LaTX samples (precursor δ-LIT and mature α-LCT) in an indirect way, since a complete LaTX tetramer is the prerequisite of functional pore-formation and thus enabling measurements of single channel currents. Indeed, our samples showed full activity and ion channel gating events of single pore units were detected under the conditions of the high resolution electrophysiological experiment. This is in complete agreement with previous experiments on α-LTX purified from the venom, which was also shown to form pores on artificial lipid bilayers, but efficient incorporation into biological membranes was only achieved in presence of specific receptors^17–19^. With regard however to our experiments on δ-LIT, we assume that rapid concentration of the sample during cryoEM grid preparation might have been an important factor, for successful assembly of the soluble tetramer, even for a small subpopulation of particles. The charge on the artificial bilayers might be another important factor for efficient LaTX recruitment and subsequent pore formation in the electrophysiology experiments, due to the bipolar charge distribution of the AR-domains. Under physiological conditions, the individual LaTX receptors are however apparently the critical factors for efficient toxin recruitment, assembly of the tetramer and subsequent pore formation^17–19^. Similar dependencies are well known for prepore oligomers of other toxins assembling at the membrane prior pore formation^52^.Oligomerization might also be reinforced by additional factors in the venom of the spider or receptor mediated interactions at the outer cell surface. Latrodectins, low molecular weight proteins characterized from the black widow venom, are known for example to associate to LaTXs and suspected to enhance their potency by altering the local ion balance^53^.

### Implications for latrotoxin receptor recognition, pore-formation and Ca^2+^ sensing

Neurexin and latrophilin are two well-studied receptors of latrotoxin^36,54^, but the receptor binding site (also for their respective invertebrate homologues) on LaTXs is still unknown. On the one hand, the helical bundle, connector and plug domains are buried deeply in the “crane claw”-like tetramer and the empty space below the helical bundles is most probably required for subsequent pore formation events (**Fig. 5a**). The four outer “claw fingers” formed by AR-domains contribute on the other hand the largest exposed surface for receptor interaction and indeed, ankyrin-like repeat domain are widely involved in maintaining protein-protein interactions, as the large numbers of modular repeats and adaptive surface residues make the motif a versatile protein binding partner^55,56^. In addition, our data suggest, that the inhibitory domain located at the tail of the AR-domain, probably interrupts the toxin-receptor interface. Therefore, LaTX uses most likely the lower half ARs for receptor recognition. Different LaTXs vary indeed in the number of their ARs, but high resolution structures of LaTX-receptor complexes are now necessary, towards understanding their specificity in detail.

**Figure 5.**
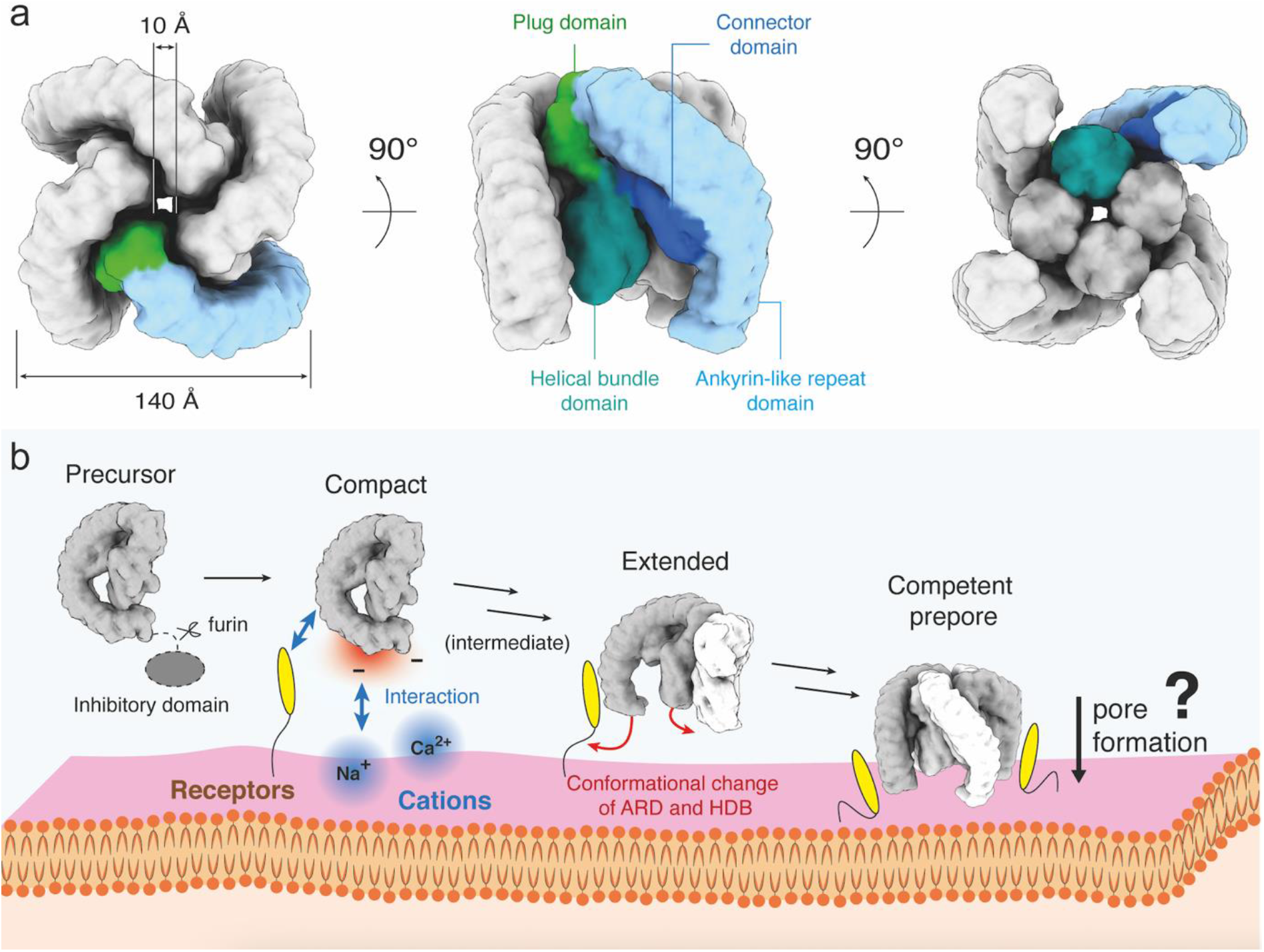
Latrotoxin tetramer assembly prior membrane insertion. **a**, Simulated volume of a soluble **δ-LIT** insertion-competent tetramer. **b**, Model of latrotoxin action at the presynaptic membrane.

Interestingly, the helical bundle domain of LaTX is reminiscent of domain I of the pore forming Cry toxin, in which six (but not five) amphipathic helices surround a hydrophobic central helix. Moreover, an α-helix in Cry toxin domain I is also interrupted by a short loop^57,58^as we observed in the H7 and H9 helices in our structure. Although the exact pore formation mechanism of Cry toxins is yet unknown, an ‘umbrella’ model of toxin-insertion, derived from structural studies on colicin, is widely accepted^59^. Recently, the RhopH complex, a pore forming protein of malaria parasites was in addition shown to possess an intriguing similar helical bundle domain to the helical bundle domain of LaTX^60^. This suggests a common strategy in until recently unrelated pore forming proteins, to protect central hydrophobic surface helices prior membrane insertion, but further studies are now required to unravel possible similarities in the respective pore formation mechanisms.

It is known that LaTX pores are permeable for cations and small molecules such as ATP or acetylcholine. The present electrophysiological measurement further reveal conductance values that in the order of magnitude typically observed for, than K^+^or Ca^2+^specific channels^61,62^. The LaTX pore is stabilized by Ca^2+^and has a preferred permeability for this ion, suggesting binding sites in the LaTX pore specialized for Ca^2+^sensing. A flexible loop in the pore can match both requirements: four loops might form an ion filter-like structure through coordinate bonds with a Ca^2+^, which can further facilitate the intake of the following Ca^2+^. In the absence of Ca^2+^, the loops become disordered, as observed for KcsA in the absence of K^+^-ions in the filter region^63^, but the pore might be then large enough to pass through the other substrates. Aspartic acid (Asp) and glutamic acid (Glu) residues have been known to play a critical role in Ca^2+^ filtering^62,64^. There are three Asp residues and five Glu residues strictly conserved at the helical bundle domain: Asp232, Asp289, Asp427, Glu121, Glu132, Glu146, Glu185, and Glu203 (residue numbers according to δ-LIT).

### LaTX mechanism of oligomerization prior membrane insertion

Taking all observations together, we propose a four-step mechanism of oligomerization and membrane binding of LaTXs (**Figure 5b**). Firstly, the inhibitory domain of LaTX is removed by proteolytic cleavage. This enables the toxins (“compact” and flexible intermediate conformations”) to be recognized by receptors at the extracellular side of the cell membrane. The negatively charged C-terminal tails of AR-domains are further attracted by the cations (e.g. Na^+^or Ca^2+^) at the extracellular side of the cell membrane, which might be crucial to orientate the molecules properly, with the “claw tips” directly facing the membrane. Adjacent protomers plug to each other via their AR-domains and plug domains in a cyclic sequential manner towards the formation of the tetramer. For α-LCT (and possibly also for other LaTXs with 22 ARs), this interaction is expected in addition to trigger a conformational change for each protomer and stabilize an extended conformation. The resulting tetramer resembles in shape an open “crane claw” (prepore; membrane insertion competent state). Notably, in this orientation, the bottom part of the cylindrical helical bundle domains, composed each of five parallel aligned helices protecting the central putative transmembrane helix, are exposed, perpendicular aligned and directly facing the membrane. This orientation towards the membrane, suggests that subsequent pore formation events, might involve massive rearrangements within the four helical bundle domains, resulting, among other events, into downwards injection of the shielded H8 helices for synchronized membrane penetration, oligomerization in the membrane and finally formation of a transmembrane channel.

## Concluding remarks

Our cryoEM results reveal the general architecture of LaTXs and allow us in combination with first functional studies to understand key steps of LaTX action at molecular level. Future studies of receptor-bound LaTXs in membrane inserted state together with mutational analysis of Ca^2+^sensing candidate loops and subsequent electrophysiological studies will be necessary to shed light into the intriguing structure and physiological function of the LaTX pore.

## Acknowledgments

We thank Dr. O. Hofnagel and Dr. D. Prumbaum for assistance with dataset acquisition and the development team of the SPHIRE software suite for assistance with image processing. This work was supported by funds from Uehara Memorial Foundation Overseas Postdoctoral Fellowships, Japan Society for the Promotion of Science (JSPS) Overseas Research Fellowships and the Humboldt Research Fellowship for Postdoctoral Researchers (to M.C.), the Max Planck Society (to S.R.) and the Medical Faculty of the University of Münster (to C.G.).

## Author Contributions

C.G. conceived and supervised the study, M.C. designed latrotoxin expression and purification experiments, M.C., L.E. purified samples, performed cryoEM experiments and processed cryoEM data; M.C. analyzed cryoEM data, performed model building and interpreted data with contributions from S.R. and C.G.; D.B. performed electrophysiological analysis. R.W., D.B. analyzed electrophysiological data, M.C. drafted the manuscript, C.G., R.W. wrote the manuscript, S.R., R.W., C.G. revised the manuscript. All authors read and approved the final manuscript.

## Data Availability

The cryoEM maps of α-LCT monomer and δ-LIT dimer have been deposited to the Electron Microscopy Data Bank (EMDB) under the accession codes XXX and XXX. The respective cryoEM datasets have been deposited to EMPIAR under accession codes XXX and XXX. The coordinates of the corresponding models have been deposited to the Protein Data Bank (PDB) under accession codes XXX and XXX. Other data are available from the corresponding author upon reasonable request.

