## Supplementary Information for "Molecular architecture of black widow spider neurotoxins"

# of

**Contents:**

1. Supplementary Figures
2. Supplementary Tables
3. Supplementary Movie Legends
4. Methods

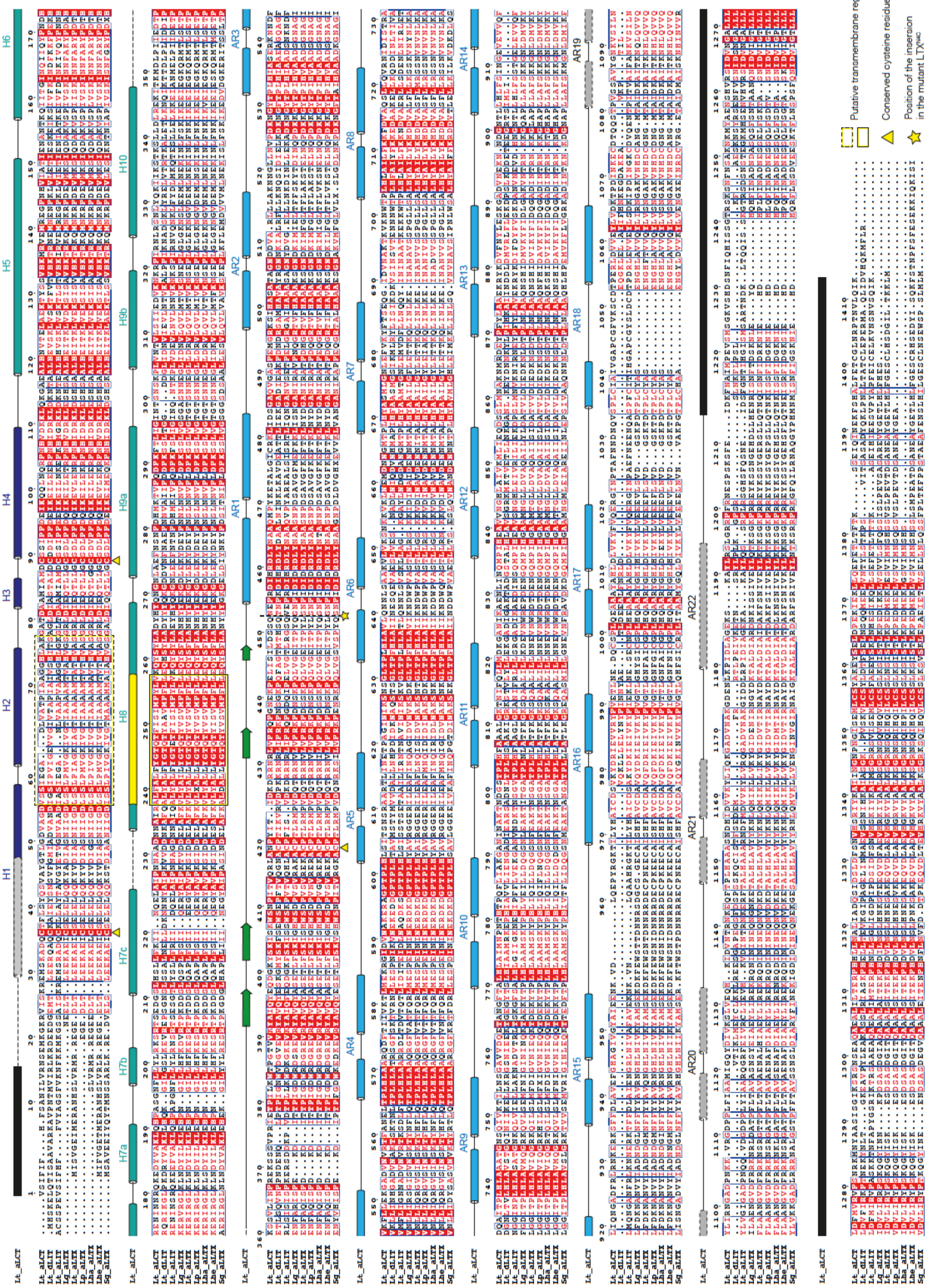
Putative transmembrane region
Conserved cysteine residues
Position of the insertion in the mutant LTX

**Supplementary Figure 1**. **Sequence alignment of latrotoxin protein sequences.** Lt: *Latrodectus tredecimguttatus* (Mediterranean black widow), Lg: *Latrodectus geometricus* (brown widow), Lp: *Latrodectus pallidus* (white widow spider), Lha: *Latrodectus hasselti* (redback spider), Lhe: *Latrodectus hesperus* (western black widow), Sg: *Steatoda grossa* (brown house spider). The secondary structure of α-LCT is shown at the top. Dashed lines or black boxes indicate disorder and/or flexible regions not included in the α-LCT model.

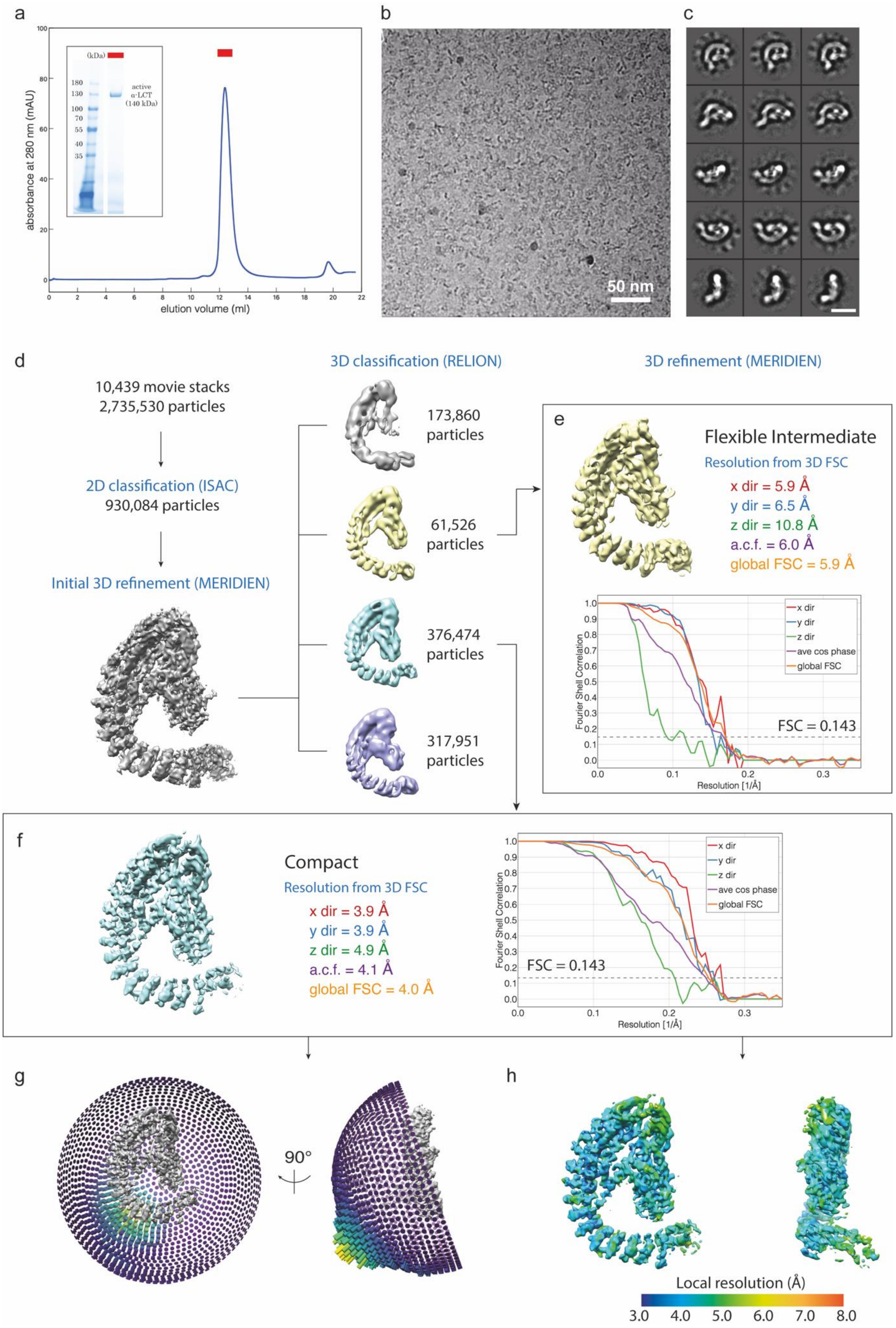

a
absorbance at 280 nm (mAU)
elution volume (ml)
(kDa)
active
α-LCT
(140 kDa)
b
50 nm
c
d
10,439 movie stacks
2,735,530 particles
2D classification (ISAC)
930,084 particles
Initial 3D refinement (MERIDIEN)
3D classification (RELION)
173,860 particles
61,526 particles
376,474 particles
317,951 particles
3D refinement (MERIDIEN)
e
Flexible Intermediate
Resolution from 3D FSC
x dir = 5.9 Å
y dir = 6.5 Å
z dir = 10.8 Å
a.c.f. = 6.0 Å
global FSC = 5.9 Å
Fourier Shell Correlation
Resolution [1/Å]
FSC = 0.143
x dir
y dir
z dir
ave cos phase
global FSC
f
Compact
Resolution from 3D FSC
x dir = 3.9 Å
y dir = 3.9 Å
z dir = 4.9 Å
a.c.f. = 4.1 Å
global FSC = 4.0 Å
Fourier Shell Correlation
Resolution [1/Å]
FSC = 0.143
x dir
y dir
z dir
ave cos phase
global FSC
g
90°
h
Local resolution (Å)
3.0 4.0 5.0 6.0 7.0 8.0

25
26

**Supplementary Figure 2**. **CryoEM sample preparation, image acquisition, and data processing of α-LCT.** **a**, Size exclusion chromatography (SEC) profile of α-LCT, showing only one peak corresponding to the monomer. The insert shows an SDS-PAGE of the peak (red bar). **b**, A representative cryoEM micrograph of α-LCT. **c**, Representative reference-free 2D class averages. Scale bar: 10 nm **d**, Flowchart of α-LCT cryoEM data processing workflow. See ‘data processing’ section in Methods for details. **e, f**, 3D FSCs and final unprocessed EM maps of a flexible intermediate (e) and the best resolved compact conformation (f) of α-LCT monomer, respectively. **g**, Angular distribution for the final reconstruction of “compact” α-LCT. **h**, EM map of “compact” α-LCT colored by local resolution.

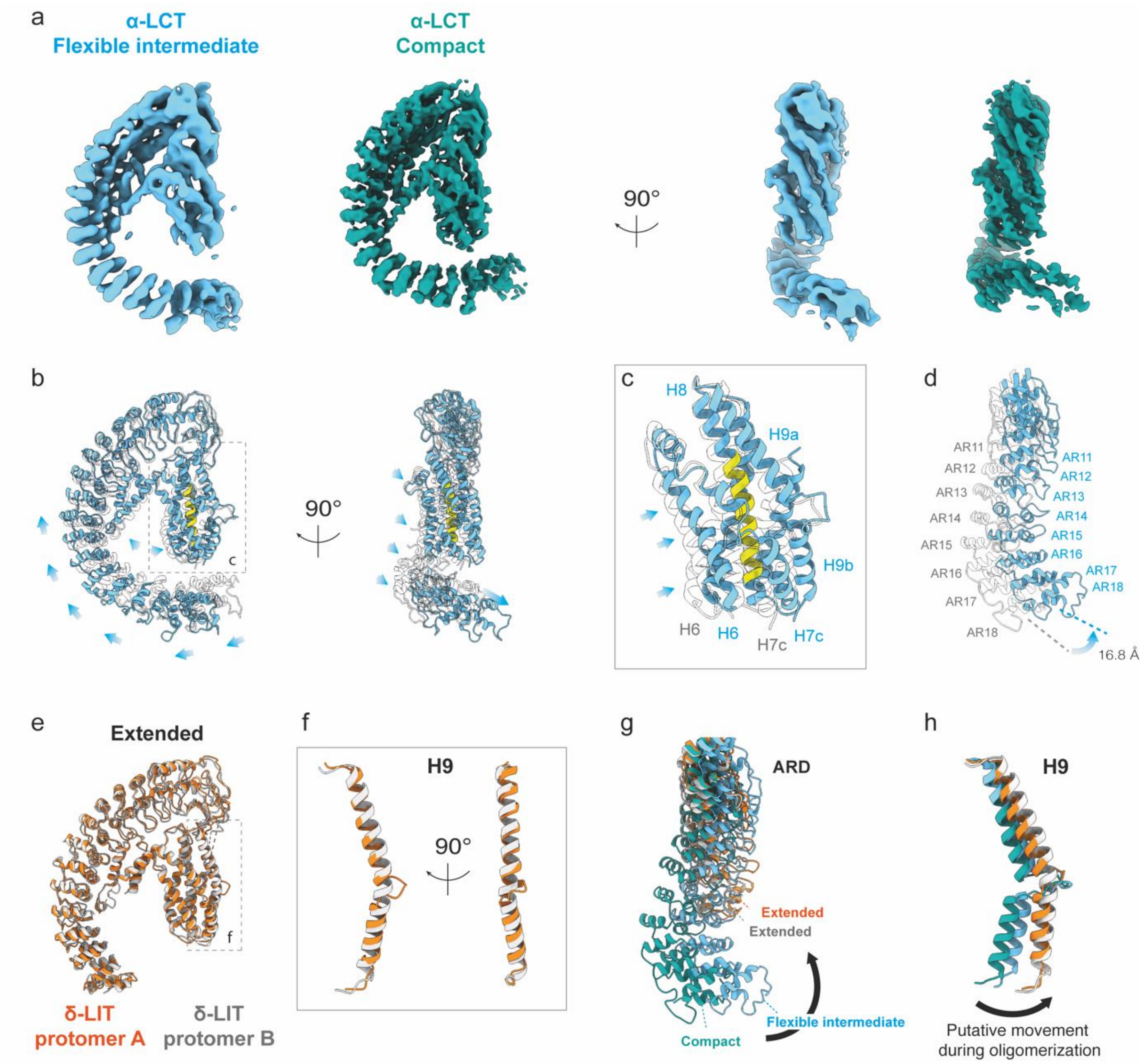

**Supplementary Figure 3**. **Different conformations of LaTX protomers. a**, Side by side comparison of the EM maps of the α-LCT monomer in the intermediate (cyan) and compact conformation (sea green). **b**, Molecular model of the α-LCT intermediate superimposed with the compact conformation (transparent). Arrows indicate domain movement from the compact to the flexible intermediate intermediate conformation. **c**, **d**, Magnified view of the helical bundle domains (c) and the terminal ARs (d) 11-18. **e**, Comparison of the two protomers of the δ-LIT dimer. Protomer A: orange; Protomer B: gray. **f**, Magnified view of the helix H9. **g,h**, Comparison of the AR-domains (ARD) and helix H9 of all four conformations, respectively.

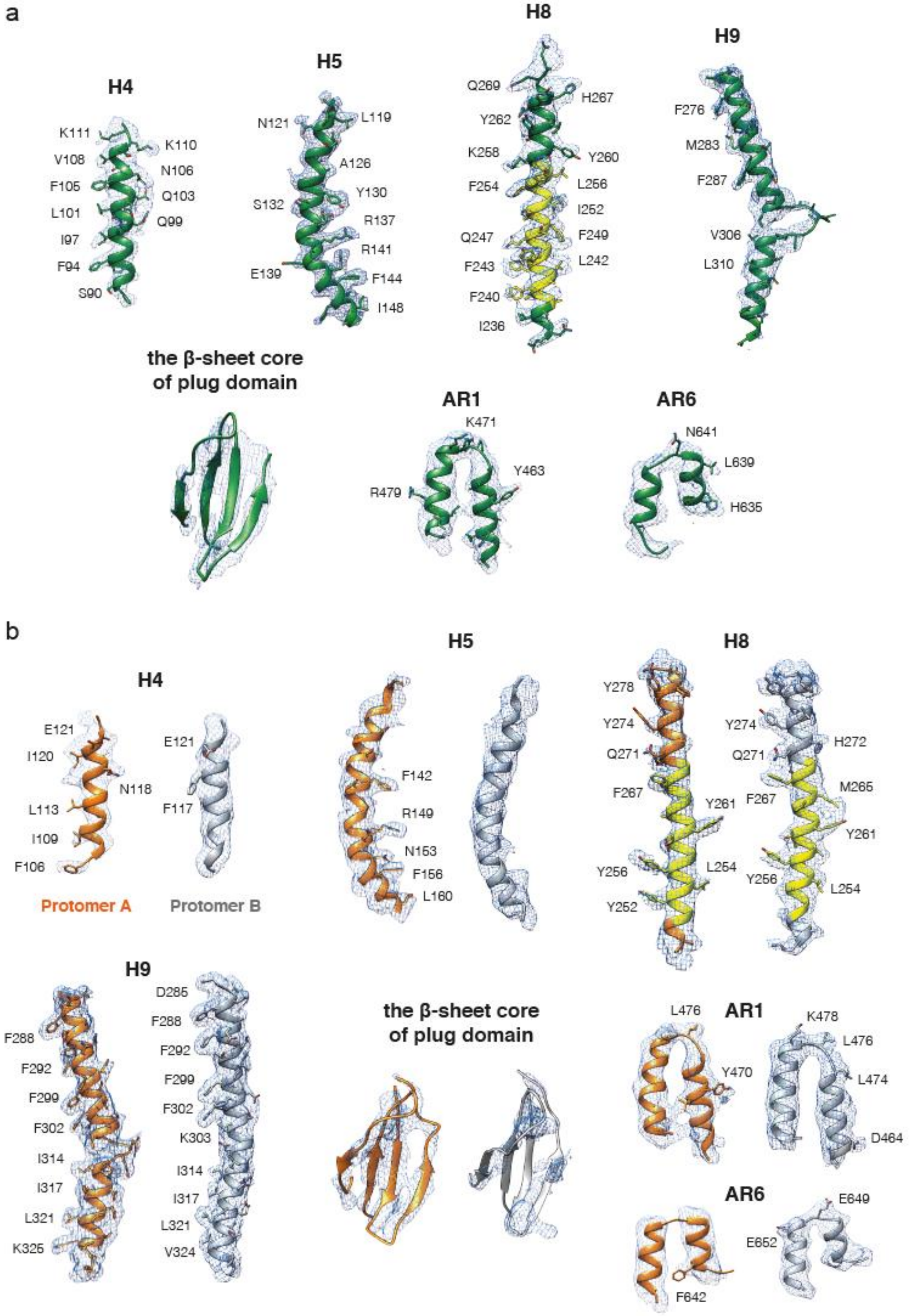

**Supplementary Figure 4. CryoEM densities of α-LCT and δ-LIT.** Superposition of segments of the molecular model of α-LCT(a) and δ-LIT (b) with the cryoEM density (mesh).

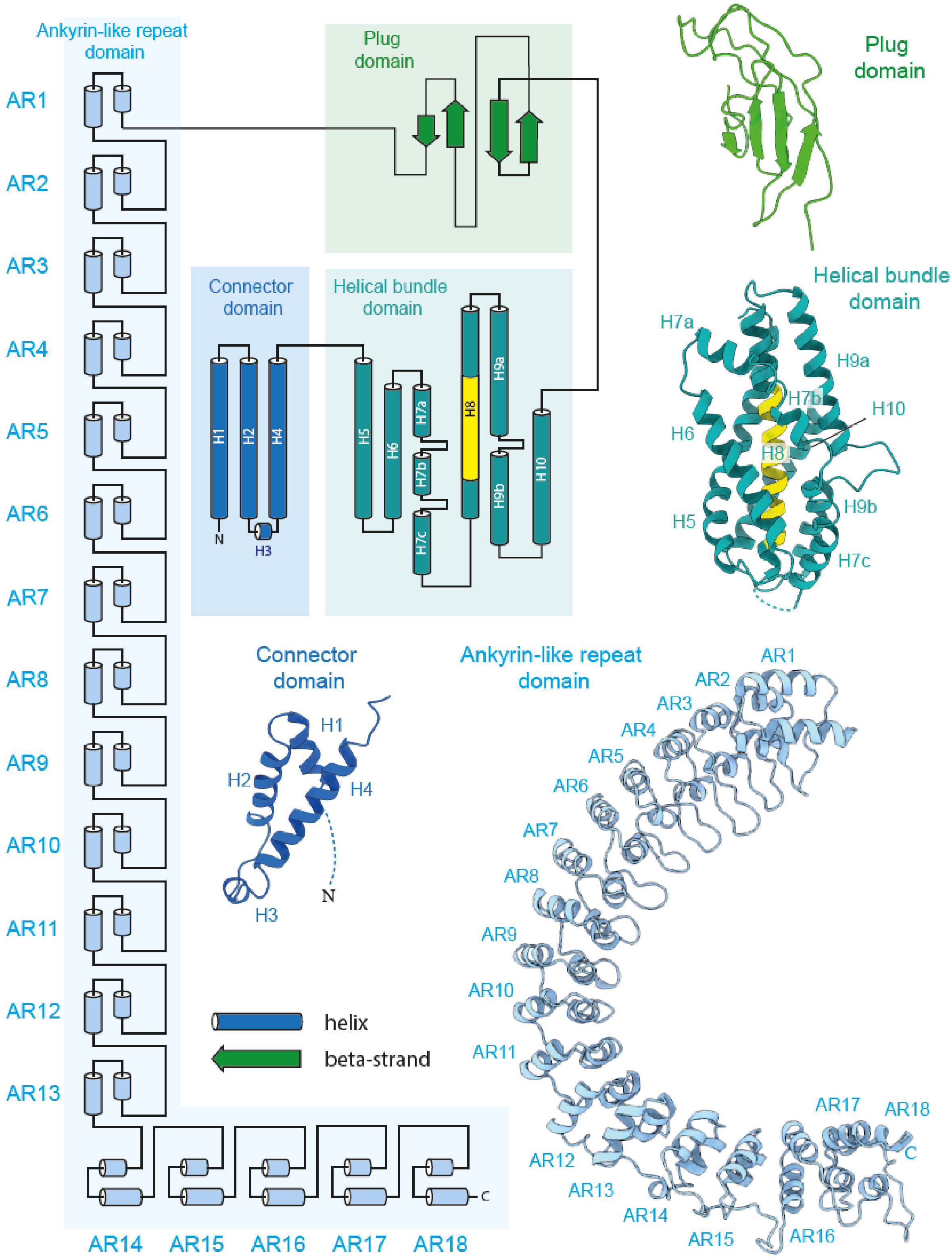

**Supplementary Figure 5**. **Topology diagram of α-LCT.** The connector domain (blue), helical bundle domain (sea green), transmembrane region (yellow), plug domain (green) and ankyrin-like repeat domain (cyan) are shown together with the respective molecular models.

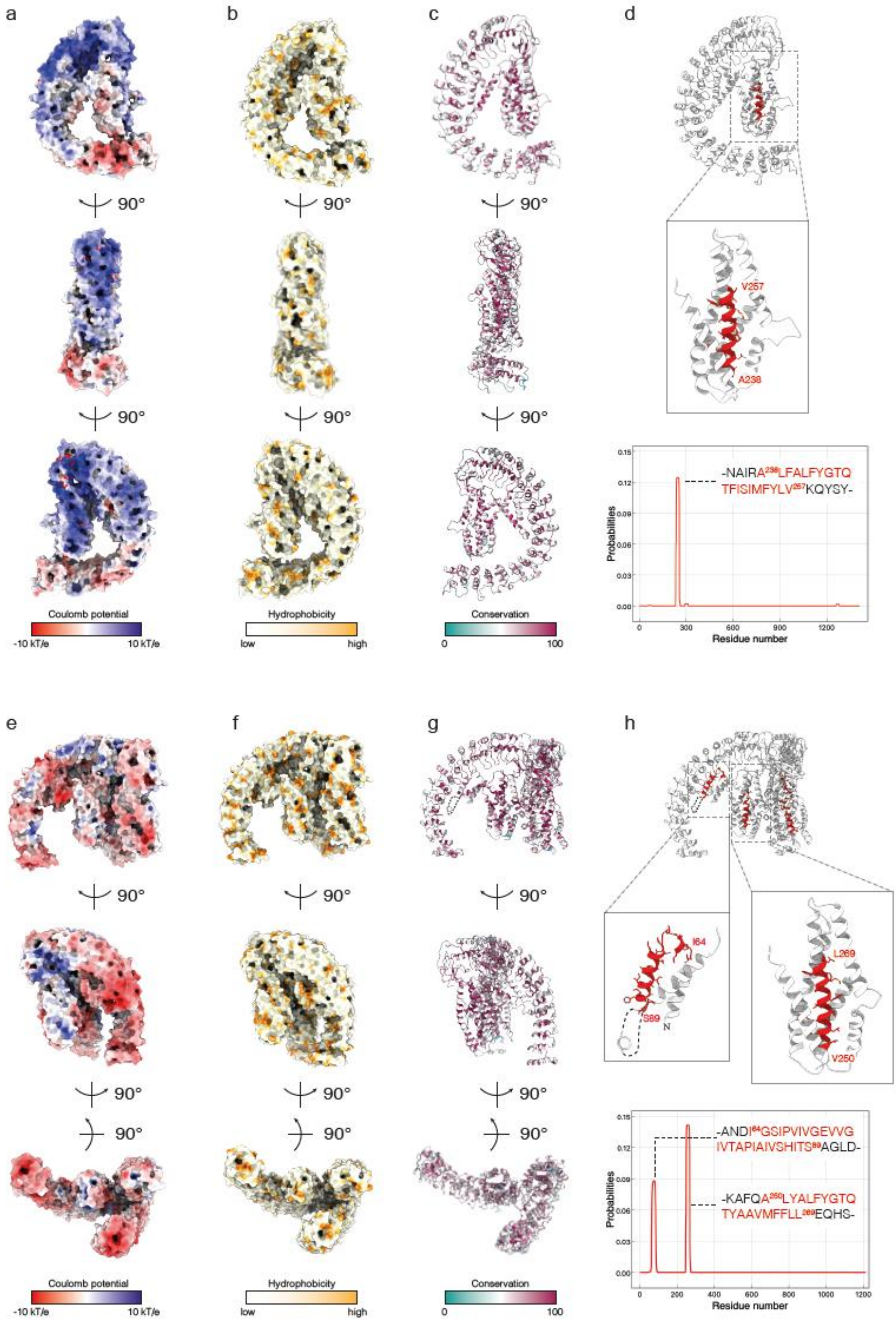
a
b
c
d
90°
V257
A238
Coulomb potential
-10 kT/e
10 kT/e
Hydrophobicity
low
high
Conservation
0
100
-NAIRA238LFALFYGTQ
TFISIMFYLV257KQYSY-
Probabilities
Residue number
e
f
g
h
I64
S89
N
L269
V250
-ANDI64GSIPVIVGEVVG
IVTAPIAIVSHITS89AGLD-
-KAFQA250LYALFYGTQ
TYAAVMFFLL269EQHS-

**Supplementary Figure 6**. **Comparative analysis of the biophysical properties and topology of the predicted transmembrane helices a**, Surface electrostatics of the α-LCT monomer at pH 7.2 calculated in APBS. Red: -10 kT/e; Blue: +10 kT/e. **b**, Surface hydrophobicity. White: low; Yellow: high **c**, Amino acid conservation. Maroon: fully conserved positions; Cyan: no conservation. **d**, The putative transmembrane region (red) of α-LCT predicted by TMHMM Server v. 2.0. **e-h**, The same structural analysis performed for the δ-LIT protomer extracted from the molecular model of the dimer. Notice there are two transmembrane regions predicted from the δ-LIT sequence.

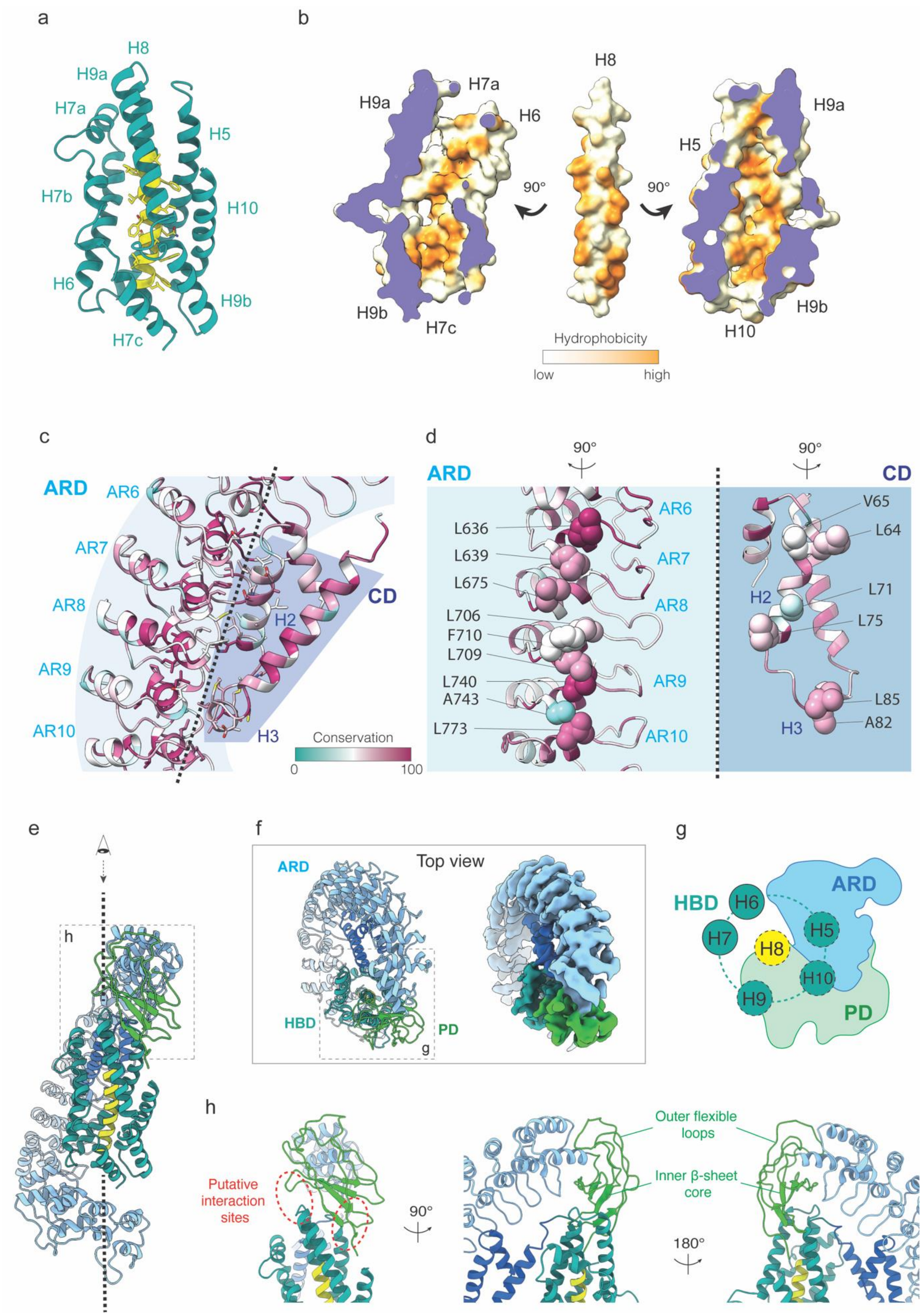
a
H8
H9a
H7a
H5
H7b
H10
H6
H9b
H7c
b
H8
H7a
H9a
H6
H9a
H5
90°
90°
H9b
H7c
H9b
H10
Hydrophobicity
low
high
c
ARD
AR6
AR7
AR8
AR9
AR10
CD
H2
H3
Conservation
0
100
d
ARD
90°
90°
CD
L636
L639
L675
L706
F710
L709
L740
A743
L773
AR6
AR7
AR8
AR9
AR10
V65
L64
L71
H2
L75
L85
H3
A82
e
h
f
ARD
Top view
HBD
PD
g
g
HBD
H6
ARD
H7
H5
H8
H10
H9
PD
h
Outer flexible loops
Putative interaction sites
90°
Inner β-sheet core
180°

**Supplementary Figure 7**. **Structural characteristicss of α-LCT a,** Close-up view of the HBD. Helix H9 is shown in the front. **b**, The hydrophobic pocket formed by the HDB. The center helix H8 is shown in the same orientation as **a**, together with sliced views of the pocket formed by the outer helices of the bundle, displaying the hydrophobic surface potential. White: low; Yellow: high. **c,** Close-up view of the CD-ARD interface. Ribbons are colored by conservation. **d**, Side views of the interface between ARD and CD. ARD and CD are rotated 90 degrees to the left and right, respectively. The residues which are considered to contribute to the hydrophobic interaction are shown as spheres. **e,f**, Side (e) and top (f) view of α-LCT monomer. The center helix (H8) of HBD in (e) is aligned to the vertical axis. **g**, A schematic diagram of the HBD-ARD-PD interface. The upper side of HBD is partially covered by the PD and ARD. **h**, Close-up view of the PD. The putative interaction sites with HBD (left) and the overall two-layer structure of PD , i.e., the flexible loops and the β-sheet core, are indicated. PD:plug domain; HBD:helical bundle domain; CD: connector domain; ARD: ankyrin-like repeat domain

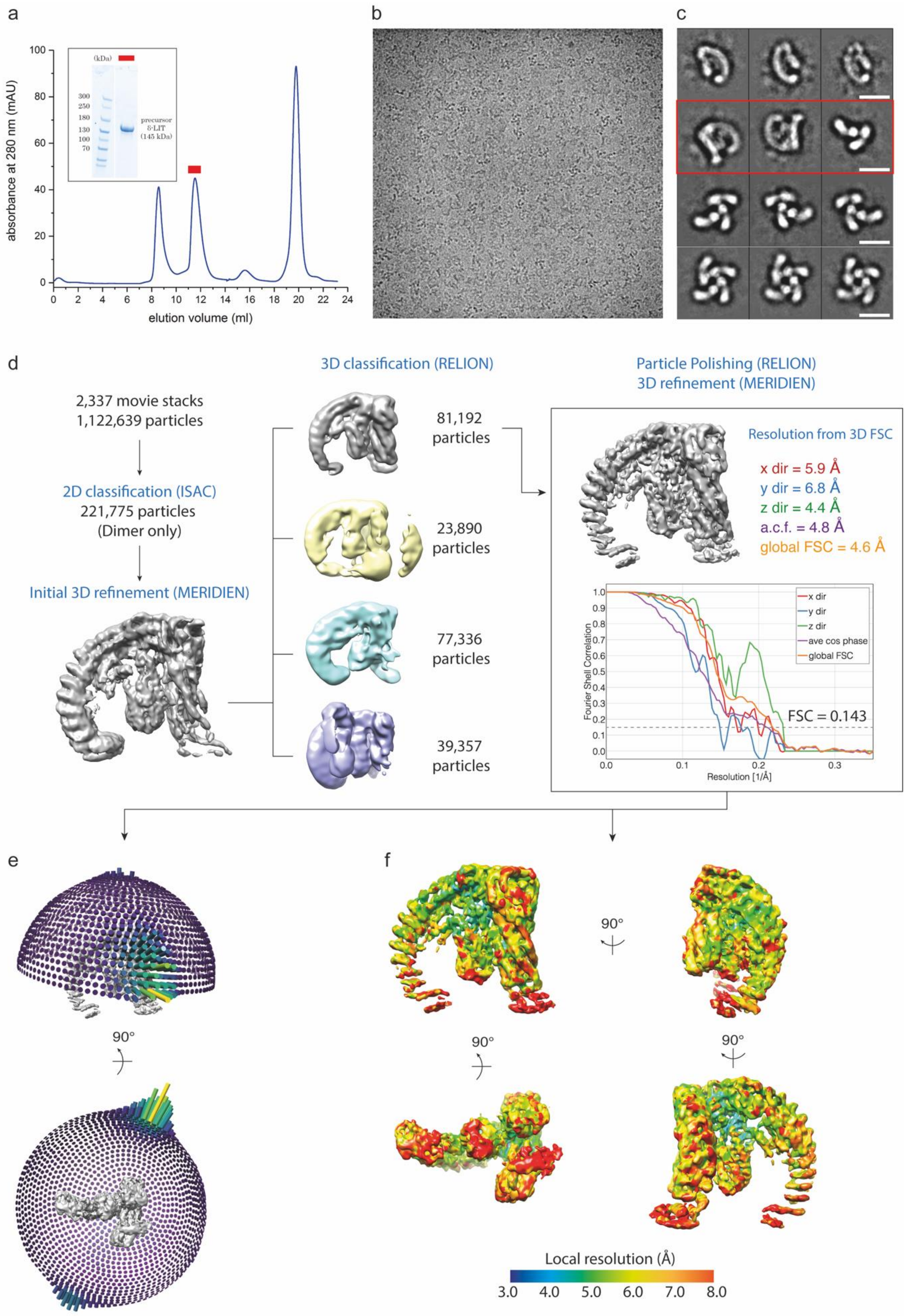
a
b
c
absorbance at 280 nm (mAU)
elution volume (ml)
(kDa)
300
250
180
130
100
70
precursor δ-LIT (145 kDa)
d
2,337 movie stacks
1,122,639 particles
2D classification (ISAC)
221,775 particles
(Dimer only)
Initial 3D refinement (MERIDIEN)
3D classification (RELION)
81,192 particles
23,890 particles
77,336 particles
39,357 particles
Particle Polishing (RELION)
3D refinement (MERIDIEN)
Resolution from 3D FSC
x dir = 5.9 Å
y dir = 6.8 Å
z dir = 4.4 Å
a.c.f. = 4.8 Å
global FSC = 4.6 Å
x dir
y dir
z dir
ave cos phase
global FSC
Fourier Shell Correlation
FSC = 0.143
Resolution [1/Å]
e
90°
f
90°
90°
90°
Local resolution (Å)
3.0
4.0
5.0
6.0
7.0
8.0

**Supplementary Figure 8. Sample preparation, image acquisition, and data processing of δ-LIT dimer.** **a**, Size exclusion chromatography (SEC) profile of δ-LIT. The dimer peak is indicated (red bar). The insert shows an SDS-PAGE of the respective peak. **b**, A representative cryoEM micrograph of δ-LIT. **c**, Representative reference free 2D class averages **d**, Flowchart of δ-LIT cryoEM data processing workflow. See 'data processing' section in Methods for details. **e, f**, 3D FSC and final unprocessed EM map of δ-LIT dimer. **g**, Angular distribution for the final reconstruction of the δ-LIT dimer **h**, The EM map of δ-LIT colored by local resolution.

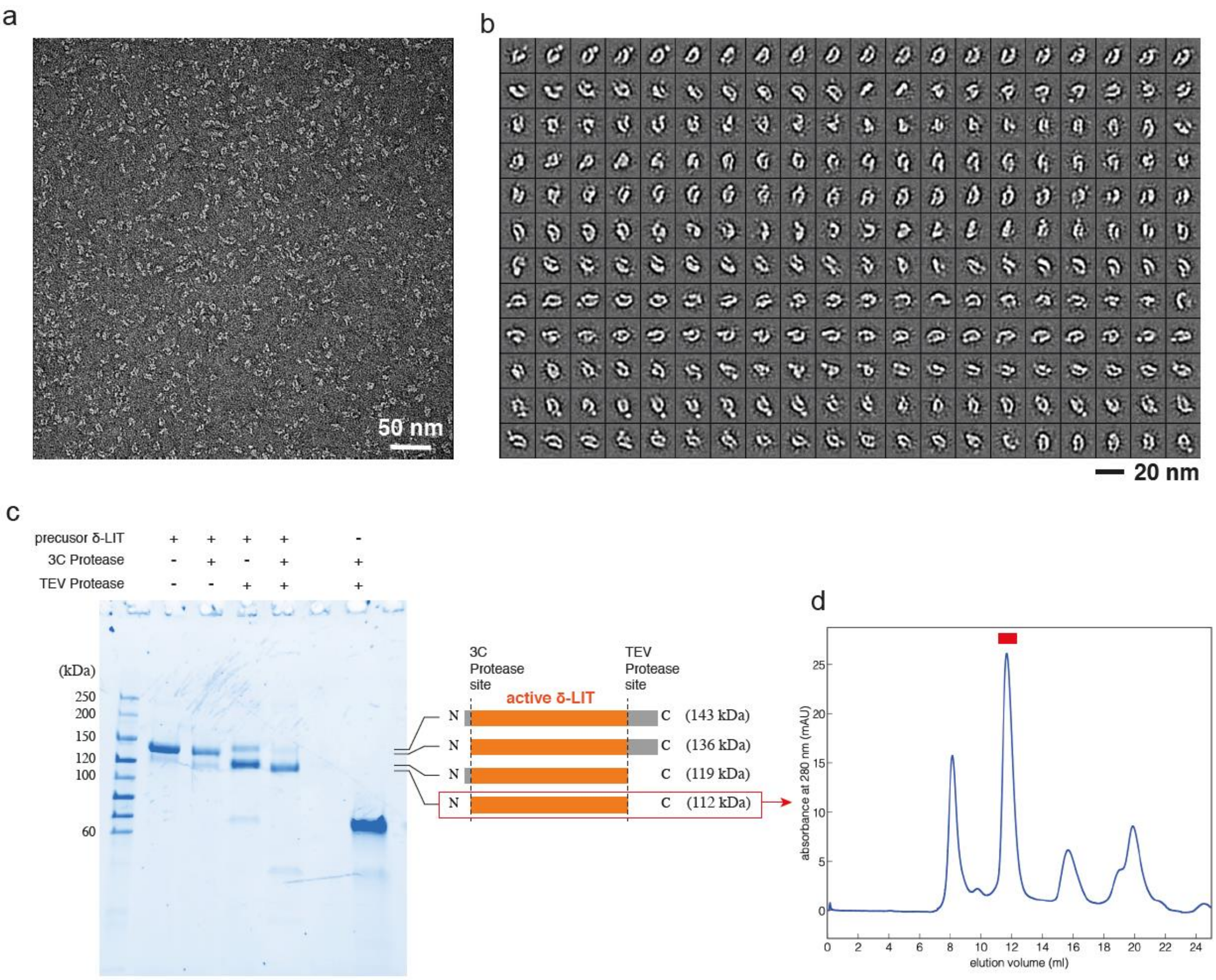

**Supplementary Figure 9. Negative stain-EM analysis of precursor δ-LIT and preparation of mature δ-LIT a**, Representative negative stain-EM micrograph of δ-LIT. **b**, Representative reference free 2D class averages **c**, SDS-PAGE of the δ-LIT sample after activation by proteolytic cleavage (lane 5) with negative controls. The expected primary structures and molecular weights are indicated. **d**, SEC profile of the sample after proteolytic cleavage. The third peak (red bar) was pooled and used for the following electrophysiologic analysis.

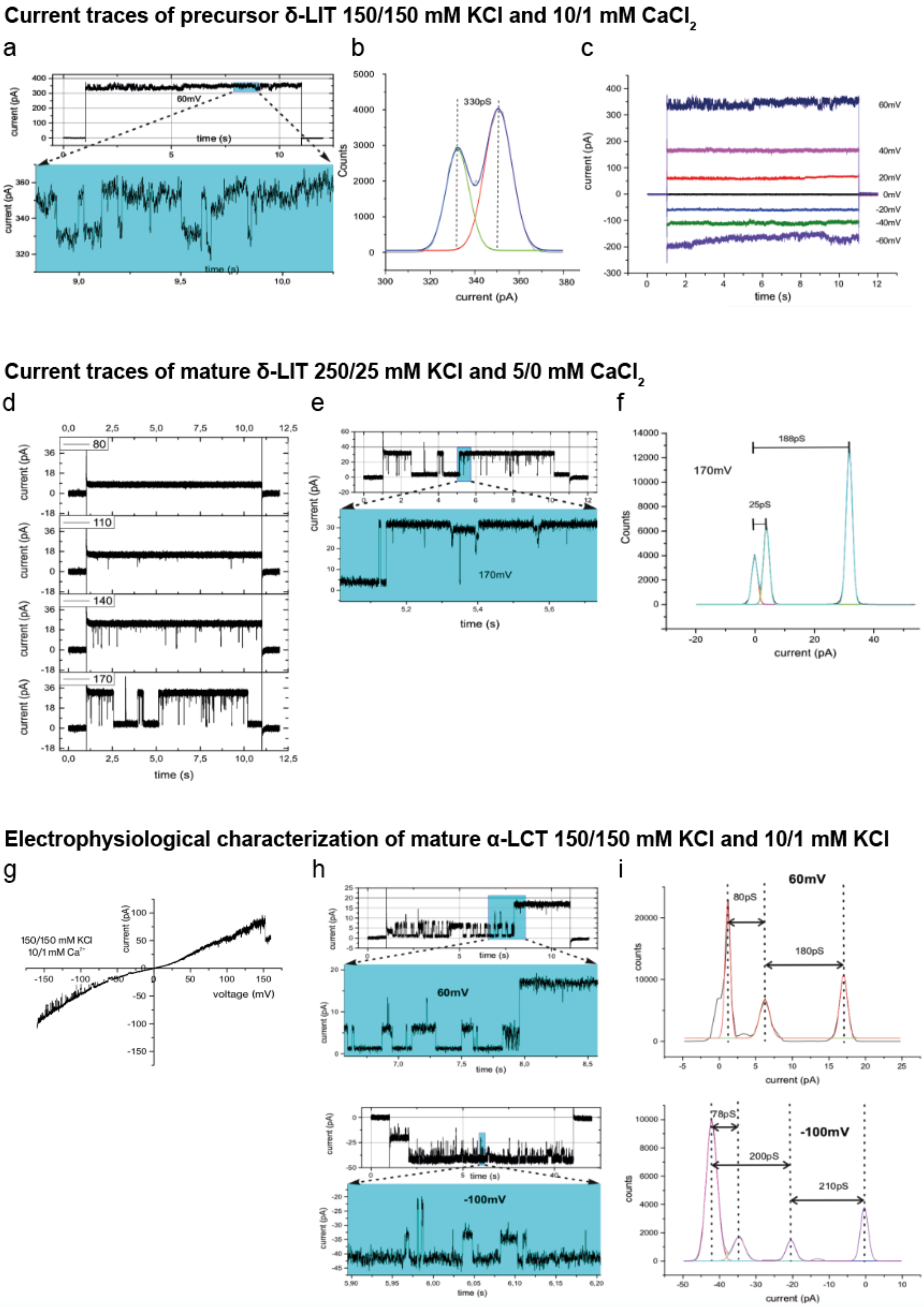
Current traces of precursor δ-LIT 150/150 mM KCl and 10/1 mM CaCl2
a
b
c
current (pA)
time (s)
60mV
330pS
Counts
current (pA)
60mV
40mV
20mV
0mV
-20mV
-40mV
-60mV
Current traces of mature δ-LIT 250/25 mM KCl and 5/0 mM CaCl2
d
e
f
80
110
140
170
current (pA)
time (s)
170mV
188pS
25pS
Counts
current (pA)
Electrophysiological characterization of mature α-LCT 150/150 mM KCl and 10/1 mM KCl
g
h
i
150/150 mM KCl
10/1 mM Ca2+
current (pA)
voltage (mV)
60mV
-100mV
time (s)
80pS
180pS
78pS
200pS
210pS
counts
current (pA)

**Supplementary Figure 10**. **Electrophysiological analysis of LaTXs. a,** Current recording and expansion plot from a bilayer containing the precursor δ-LIT in symmetrical 150/150 mM KCl (cis/trans) and 10/1 mM $CaCl_2$ buffer (cis/trans) in response to a voltage gate with $V_{cmd}$=60mV. **b,** All point current amplitude histogram from the record in (**a**). **c**, Current recordings from a bilayer containing the precursor δ-LIT in symmetrical 150/150 mM KCl and 10/1 mM $CaCl_2$ buffer (cis/trans) in response to voltage gates with the indicated amplitudes. Remarkably precursor δ-LIT channel gating was rather unstructured and noisy with fast current transitions which could not be clearly resolved in amplitude or time (**a,c**). Using a simplified cylindrical model this $\bar{G}_{main}$ would correspond to a pore restriction diameter of about 1.5 nm[1,2] **(c) d**, Current recording from a bilayer containing the reconstituted mature δ-LIT in asymmetric 250/25 mM KCl and 5/0 mM $CaCl_2$ (cis/trans) buffer at the indicated $V_{cmd}$. Like the precursor δ-LIT also the mature variant harbors a high lipid bilayer insertion activity. For single insertion analysis here the 1:120000 diluted (0.08 nM) δ-LIT finally showed a single insertion event and at all applied $V_{cmd}$ the current traces displayed at the given time-resolution clearly defined low noise gating patterns (**d,e**) **e**, Extension plot of the recording at $V_{cmd}$=170 mV. **f**, All point current amplitude histogram at $V_{cmd}$=170 mV of the respective recording in **d. g,** Current-voltage ramp recording from a bilayer containing the mature α-LCT in 150/150 mM KCl and 10/1 mM $CaCl_2$ (cis/trans) buffer revealing a slightly rectifying shape. **h**, Current recording and extension plot in response to voltage gates of $V_{cmd}$= 60 mV and $V_{cmd}$=-100 mV. **i,** Corresponding all point current amplitude histogram at $V_{cmd}$= 60mV and -100mV revealing two open channel conductance states.

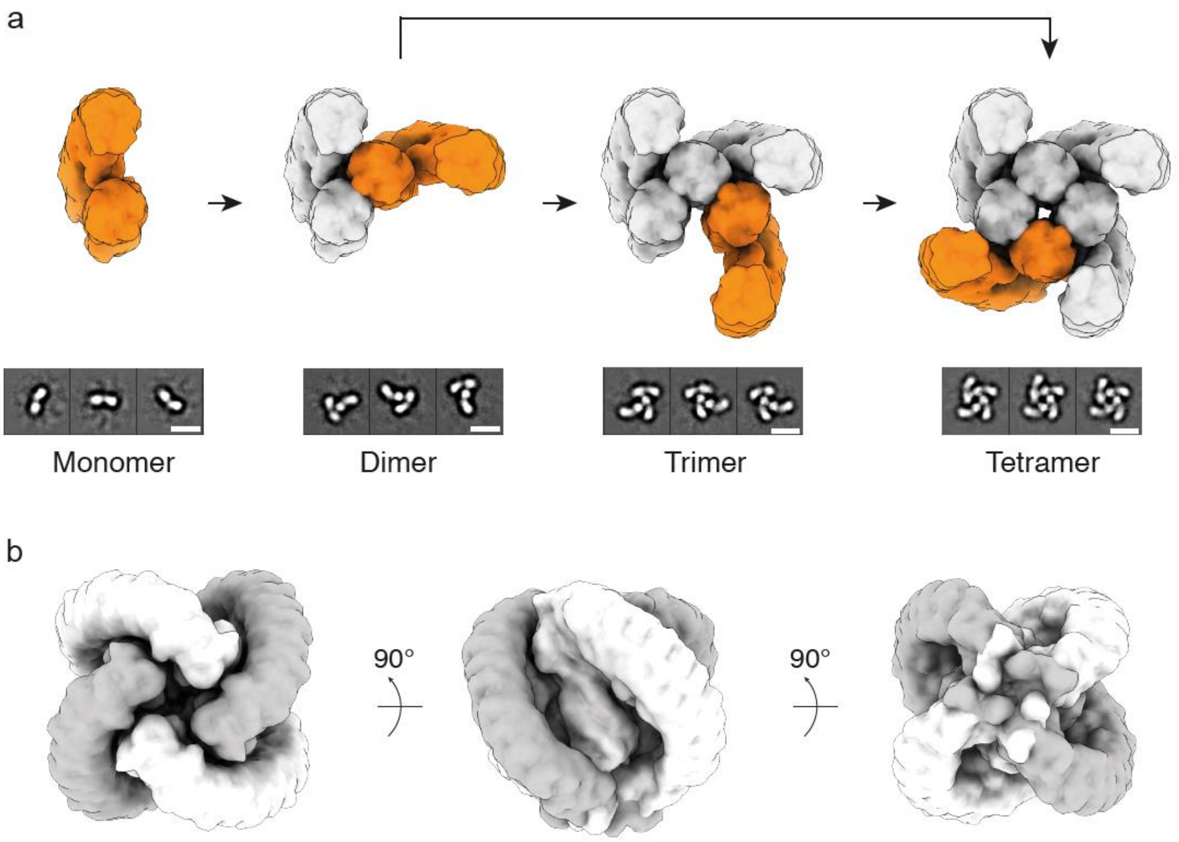

**Supplementary Figure 11. a,** Proposed sequential circular mechanism of tetramer formation of latrotoxin vie 1/2/3 mers. Representative 2D classes of each oligomer state from the present δ-LIT dataset are shown below the respective densities of simulated volumes filtered to 10 Å. **b**, Simulated volume of a tetramer assembled from “compact” state α-LCT monomers to demonstrate the mismatch to the 2D class averages of δ-LIT tetramers (see **a**) and the clashes between the AR domains. According to our model, conformational changes from the “compact” to “extended” state take place already during dimer formation and not after tetramer formation.

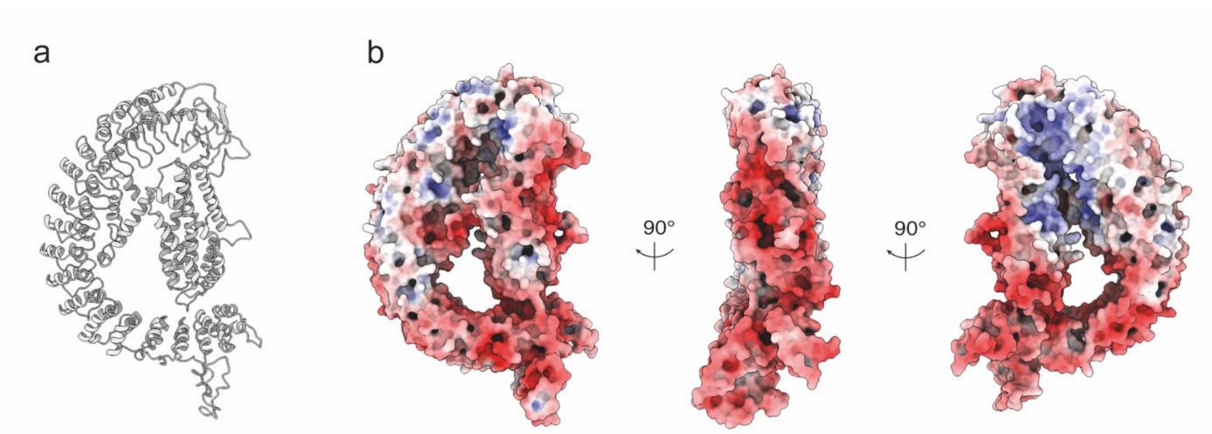

**Supplementary Figure 12.** Surface electrostatics analysis of a homology model of α-LTX **a,** A homology model of α-LTX based on the α-LCT monomer structure generated by SWISS-MODEL. **b**, Surface electrostatics of the α-LTX homology model at pH 7.2 calculated in APBS. Red: -10 kT/e; Blue: 10 kT/e.

### Protomer A

| Domain | helix | residue |
|---|---|---|
| HBD | H9 | Asp310 |
| | | Ser304 |
| | | Asp300 |
| | | ILE297 |

| | | |
|---|---|---|
| PD | | Lys435 |
| | | Lys434 |
| | | Pro433 |
| | | Val431 |
| | | Arg430 |
| | | Lys428 |
| | | Ser371 |
| | | Glu372 |
| | | Val373 |
| | | Asn374 |
| | | Phe375 |
| | | Pro376 |
| | | Asn377 |
| | | Glun381 |
| | | Val419 |
| | | Gln420 |
| | | Gly421 |

### Protomer B

| residue | helix | Domain |
|---|---|---|
| Lys184 | H6 | HBD |
| Arg188 | | |
| Leu192 | | |
| Lys200 | H7a | |
| Ala199 | | |

| | | |
|---|---|---|
| Val74 | H2 | CD |
| Glu649 | AR6 | ARD |
| Asn648 | | |
| ILE647 | | |
| Ser615 | AR5 | |
| Met614 | | |
| Phe583 | AR4 | |
| Ser581 | | |
| Arg580 | | |
| Glu546 | AR3 | |
| Ala547 | | |
| Gln549 | AR3 | |
| Arg512 | AR2 | |
| Tyr511 | | |
| ILE476 | AR1 | |
| Asn477 | | |
| Lys478 | | |
| Lys480 | | |

**Supplementary Table 1.** Residues which are considered to contribute to dimerization. The spatially adjacent residues are indicated with the same color in the table. The helices and domains where the residues are located are also indicated.

| | **α-LCT monomer** | **δ-LIT dimer** |
|---|---|---|
| Microscope | Titan Krios (X-FEG, Cs 2.7 mm) | Titan Krios (X-FEG, Cs -corrected) |
| Energy filter /Slit width (eV) | GIF Quantum / 20 | GIF Quantum / 20 |
| Voltage (kV) | 300 | 300 |
| Defocus range (um) | -1.3 to -2.5 | -1.3 to -2.4 |
| Camera | K3 super-res. | K3 super-res. |
| Pixel size (Å) | 0.45 / 0.9 | 0.45 / 0.9 |
| Total electron dose (e/Å2) | 69 | 58 |
| Exposure time (sec) | 3 | 3 |
| Frames per movie | 60 | 60 |
| Nr. of images | 10439 | 2339 |

[a] Pixel size after 2x binning used for processing

**Supplementary Table 1. Data collection statistics**.

**Supplementary Movie 1.** The cryoEM map and the overall structure of α-LCT monomer.

**Supplementary Movie 2.** The cryoEM map and the overall structure of δ-LIT dimer.

**Supplementary Movie 3**. Putative conformational change during oligomerization. The four conformations, i.e., the “compact” (sea green; α-LCT), an intermediate (cyan: α-LCT) and the “extended” state (orange: δ-LIT protomer A; gray: δ-LIT protomer B) are shown sequentially. The morphing occurs between the “compact” conformation of the α-LCT monomer and the “extended” conformation of δ-LIT protomer B. The conformational change of the overall structure is shown followed by close-up views of the helical bundle-, connector- and AR-domain

### Methods

#### Protein expression and purification

Our expression protocol was modified from Volynski K.E., et. al.[3] based on secretion in the baculovirus expression system. The cDNAs of mature α-LCT (residues 16-1211; UniProtKB Q9XZC0) and full-length δ-LIT (residues 1-1214; UniProtKB Q25338) were optimized for recombinant protein expression in insect cells and fused with an N-terminal honeybee melittin signal peptide (MKFLVNVALVFMVVYISYIY)[4] to enhance the efficiency of secretion. A Strep-tag II followed by an HRV-3C protease cleavage site was added after the melittin peptide, and a C-terminal $His_8$ was added with a thrombin cleavage site. All DNA fragments were synthesized (GenScript Biotech) and cloned into a pACEBac1 plasmid[5]. For producing the mature δ-LIT, Domain I (residues 1-28) was deleted from the construction, and a TEV protease site was inserted between Domain III (CTD) and Domain IV (after residue 1019). All the plasmids used in the present study are shown in Extended Data.

The bacmids were generated by transforming 200 ng of each plasmid to DH10EMBacY *E.coli* cells. Positive baculovirus genomes were selected using blue/white screening on LB-agar plates containing 100 μg/ml Ampicillin, 10 μg/ml Gentamicin, 10 μg/ml tetracycline, 500 μg/ml 5-Bromo-4-chloro-3-indolyl-β-D-galactopyranoside (X-Gal), and 0.5 mM Isopropyl β-D-1-thiogalactopyranoside (IPTG). The white single colonies were inoculated into 5 ml LB containing 10 μg/ml Gentamicin at 37 ℃ for 24 hours. The *E.coli* cells were lysed by the P1, P2, and N3 buffers of Plasmid Miniprep Kit (QIAGEN), followed by isopropanol precipitation. 700 μl isopropanol was applied to 800 μl cell lysate. The pellets were collected by centrifugation at 16,000g for 10 min, washed with 200 μl 70% (v/v) ethanol, and solubilized in 50 μl sterilized water. The final concentration of bacmid was approximately 3 μg/μl.

For generating the virus stocks, 10 μg of each bacmid were added with 250 μl Sf-900 II serum-free medium (Thermo Fisher Scientific) and 4 μl of FuGENE HD Transfection Reagent (Promega). The mixtures were transfected into 3 ml of $0.5 \times 10^6$ cells/ml Sf9 cells and the supernatants, namely the $V_0$ virus stocks, were collected after incubation at 27 ℃ for 72 hours. In order to obtain the virus stocks in higher titration and larger amount, $V_1$ and $V_2$ virus stocks were generated by suspension culture of the $1.0 \times 10^6$ cells/ml Sf9 cells infected with 0.2%(v/v) virus from the previous step at 27 ℃ in 100 rpm for 72 hours and stored at 4 ℃ with 10% fetal bovine serum (FBS) (Thermo Fisher Scientific). The titration of the final $V_2$ virus stocks was approximately $6.0 \times 10^8$ PFU/ml measured by plaque assay.

For large scale expression, $2.0 \times 10^6$ cells/ml Hi5 cells were infected with the $V_2$ virus at a multiplicity of infection (MOI) of 1-2. After suspension culture at 27 ℃ in 100 rpm for 72 hours, the Hi5 cells were removed by centrifugation. The supernatants containing the secreted proteins were chilled to 4 ℃ and adjusted to pH 8.0 by adding Tris. All the following procedures were conducted at 4 ℃ unless otherwise noted. White precipitation formed during pH adjustment and was removed by passing it through a 0.45 μm filter (Millipore).

Cleared supernatants were applied to gravity-flow columns filled with 10 ml of Strep-Tactin Sepharose resin (iba) and equilibrated in Wash Buffer (100 mM Tris-HCl pH 8.0, 150 mM NaCl, 1 mM Ethylenediaminetetraacetic acid (EDTA)). Subsequently, the columns were washed with 50 ml (5 CV) of Wash Buffer and eluted with Elution Buffer (100 mM Tris-HCl pH 8.0, 150 mM NaCl, 1 mM EDTA, 2.5 mM desthiobiotin). The elutes were fractionated and confirmed by SDS-PAGE. The fractions containing target proteins were pooled and concentrated to 0.5-5 ml. The protein samples were further purified by size exclusion chromatography (SEC) on Superdex 200 increase 10/300 or Superdex 200 26/60 column (GE

Healthcare) equilibrated in Wash Buffer. The final concentration was measured by the Bradford method (Bradford protein assay kit, Bio-Rad) and the purity was confirmed by SDS-PAGE and negative stain electron microscopy.

The mature δ-LIT was generated by incubating the elution of Strep-tag affinity chromatography with 10%(w/w) His-PreScission 3C protease and TEV protease (provided by the Protein Chemistry Facility, Max-Planck Institute for Molecular Physiology) at 4 ℃ overnight. The product was confirmed by SDS-PAGE followed by SEC as aforementioned.

**Negative stain-EM analysis**

Negative stain-EM was applied to all the three toxin samples to assess sample quality prior cryoEM analysis. 4 μl of each toxin sample with a concentration of 0.01 – 0.02 mg/ml were applied to a copper grid covered by a carbon layer and incubated for 2 min at room temperature. The excess protein solution was blotted with filter paper. The grids were washed with 10 μl double-distilled water and 0.75% uranyl formate once each, followed by staining with 10 μl 0.75% uranyl formate for 45 sec. Image acquisition was performed using a JEOL JEM-1400 transmission electron microscope operating at an acceleration voltage of 120 kV. Datasets were acquired with a 4k × 4k CMOS camera F416 (TVIPS) at a magnification of 80,000 x. The defocus range was approximately -0.8 μm to -1.8 μm at a pixel size of 1.3 Å/px.

**Sample vitrification and cryoEM data acquisition**

For cryoEM sample preparation, 4 μl of protein solution were applied onto a freshly glow-discharged holey carbon grid (QUANTIFOIL R 1.2/1.3 mesh 300) and vitrified using a Vitrobot cryo-plunger (Thermo Fisher Scientific). The plunging condition was optimized by several rounds of screening sessions. The final concentrations of mature α-LCT and full-length δ-LIT solution were 0.6 mg/ml and 0.4 mg/ml, respectively.

Two datasets of the α-LCT were collected with a 300 kV Titan Krios microscope (Thermo Fisher Scientific) equipped with an X-FEG and operated by the software EPU (Thermo Fisher Scientific). One image per hole with defocus of -1.3 to -2.5 μm was collected with the K3 Summit (Gatan) direct electron detector in super-resolution mode at a magnification of 105,000 x and a corresponding pixel size of 0.45 Å/px with a GIF quantum-energy filter set to a filter width of 20 eV. Image stacks with 60 frames were collected with a total exposure time of 3 seconds and a total dose of 69.1 e/Å$^2$. These two datasets were combined and used for the following processing.

Datasets of the δ-LIT were collected with a 300 kV Titan Krios microscope (Thermo Fisher Scientific) equipped with an X-FEG and a Cs corrector and operated by the software EPU (Thermo Fisher Scientific). One image per hole with defocus of -1.3 to -2.4 μm was collected with the K3 Summit (Gatan) direct electron detector in super-resolution mode at a magnification of 81,000 x and a corresponding pixel size of 0.45 Å/px with a GIF quantum-energy filter set to a filter width of 20 eV. Image stacks with 60 frames were collected with a total exposure time of 4 seconds and a total dose of 78.7 e/Å$^2$.

The details of the dataset collection are summarized in Supplementary Table 2.

**Image processing and 3D reconstruction**

Image transfer, motion correction, and CTF estimation was performed using Motioncorr2 and CTFFIND4 implemented with TranSPHIRE[6], a recently released software package that allows automated on-the-fly processing for cryoEM. In particular, the motion correction was performed by MotionCor2[7] to create aligned dose-weighted averages. The super-resolution images were binned twice after motion correction to speed up subsequent processing steps. The non-dose weighted average micrographs were used for CTF estimation performed by CTFFIND 4.1.10[8]. Outliers were removed based on the estimated defocus values and resolution limits. Further image processing was performed using the software package SPHIRE[9] unless otherwise noted.

Single particles were automatically picked by crYOLO[10] based on a manually trained model and extracted with a final window size of 224 × 224 pixels. The 2D classification was performed by ISAC[11] at a pixel size of 3.1 Å/pixel. The Beautifier tool implemented in SPHIRE was utilized for obtaining refined 2D classes. Classes displaying high resolution features were selected and combined into a subset. For the α-LCT monomer, an initial 3D model was generated by RVIPER[12] from a previously collected test dataset and subsequently used for 3D refinement in MERIDIEN[9] with C1 symmetry. Since the resulting 3D reconstruction showed anisotropic resolution and several low resolution regions, several rounds of 3D classification were performed by SPHIRE and RELION 3.1[13,14] until the 'compact' and 'flexible intermediate' classes were separated from the other conformations. The 'compact' subset containing 376,474 particles was further improved by CTF refinement in RELION 3.1, followed by a final round of 3D refinement in MERIDIEN. The 'flexible intermediate' class only contained 61,526 particles. Therefore, only standard 3D refinement was performed to generate a map for secondary structure fitting. The δ-LIT dimer dataset was processed with the same procedure as the α-LCT monomer until the 2D classification step. The window size was enlarged to 360 × 360 pixels due to the larger particle size. After sorting the monomer, dimer, trimer, and tetramer classes into subsets, a multi-reference classification of the dimer subset was performed in SPHIRE followed by an additional round of 3D classification in RELION 3.1. Eventually, a subset containing 81,192 particles was selected. The initial model for 3D refinement was generated by manually combining two α-LCT monomers, as indicated in the 2D class averages and applying a 15 Å low-pass filter. Subsequent polishing and CTF refinement were performed as described for the α-LCT dataset.

The final half-maps were combined upon masking using a 3D mask generated by the PostRefiner tool implemented in SPHIRE, which in addition automatically determines the B-factor and filters the resulting volume to estimated resolutions. The global resolutions as well as the 3DFSC were calculated at the 'gold standard' 0.143 criterion using the 3DFSC server[15]. The angular distribution and the local resolution distribution of the final map were analyzed using the o angular_distribution and sp_locres in SPHIRE

**Model building, validation, and visualization**

The *de novo* model building of the 'compact form' of the α-LCT monomer started from a 3D structure prediction using trRossetta (Yang 2020). The resulting model did not agree well with the density maps, therefore it was fragmented into several structural domains according to the secondary structure prediction from RaptorX Property[16]. Rigid body fitting of the individual domain was then performed using Pymol (The PyMOL Molecular Graphics System, Version 2.0 Schrödinger, LLC). Further refinement was conducted using the real-time molecular-dynamics simulation-based program ISOLDE[17] implemented in the visualization softwares UCSF Chimera and UCSF ChimeraX[18,19], and the resulting model was further refined using a combination of the model editing software COOT[20] and real-space refinement in Phenix[21]. The above adjustments were performed for several rounds until the model sufficiently fitinto the map. The model of the "flexible intermediate" state was generated by truncating side chains with the Chainsaw tool in CCP4[22] and fitted into the corresponding low-resolution map. The starting model of the δ-LIT dimer was generated by the homology-modeling service SWISS-MODEL[23] from the α-LCT model, which was subsequently fitted into the map using ISOLDE,

COOT, and Phenix. The disordered regions and side-chain atoms beyond Cβ in regions where side-chain density was only rarely evident were deleted during the model fitting.

To calculate the protein properties, i.e., surface electrostatics, hydrophobicities, residue conservations, and transmembrane region predictions, the sidechains were put back with ideal rotamers (Supplementary Fig. 6). The figures were prepared using UCSF Chimera and ChimeraX[18,19]. Multiple sequence alignment and the prediction of transmembrane helices were done using Clustal Omega[24] and TMHMM server v2.0[25], respectively. The surface electrostatics was calculated in APBS[26]and visualized by UCSF ChimeraX. These sequence features were illustrated by ESPript3[27] and Adobe Illustrator.

**Electrophysiology**

<u>Single Channel Recordings from Planar Lipid Bilayers and Data Analysis</u>

Planar lipid bilayer measurements using the Compact bilayer platform (Ionovation GmbH) were performed as described in detail previously[28]. In brief: if not explicitly stated otherwise, symmetric conditions (250 mM KCl, 10 mM HEPES, pH 7.0) were used in cis and trans compartments. The denomination cis and trans corresponds to the half-chambers of the bilayer unit. Reported membrane potentials are always referred to the trans compartment. Bilayer fabrication was performed on PFTE film at a 100 µm prepainted (1 % hexadecane in n-hexane) aperture with a Phosphatidylcholine (18:1) (PC) / Phosphatidylethanolamine (18:1) (PE) (7:3 ratio) lipid mixture (both lipids were purchased from Avanti Polar Lipids) in n-pentane using the "thinning method" [28]. Stock solutions of purified LaTXs with typically 0.5-2.5 mg/ml contained in a buffer of 100 mM NaCl, 20 mM Hepes, pH 7.4 were added to the cis compartment under slight stirring to a final protein concentration of ~1µg/ml to 10 ng/ml.

Ion channel currents were recorded using an EPC 10 USB amplifier (HEKA Elektronik GmbH) in combination with the Patchmaster data acquisition software (HEKA Elektronik GmbH). For data acquisition a sampling rate of 5kHz (voltage ramps) and 10 kHz (continuous recording) was used and the data were further analyzed using the Origin package (Origin Lab) and the MATLAB (MathWorks) based Ion-channel-Master software developed in our laboratory [28].

The GHK approach

The Goldman-Hodgkin-Katz approach is by far the most commonly used framework to describe ion permeability and selectivity of membranes [29,30]. Beside the principal difficulties underlying the macroscopic GHK constant field theory, which assumes independent movement of the ions through membrane pores (see references [31-33] for a detailed discussion) it has been demonstrated that the methodology can be used to obtain reliable semi-quantitative measures for permeation of charged drugs through membranes [1].

We were interested in obtaining information on the selectivity of the membrane reconstituted δ-LIT.

To characterize the ion fluxes mediated by the δ-LIT we employed the following experimental conditions for bilayer containing an unknown number of open δ-LIT channels.

**Permeability δ-LIT :** $P_{K^+} = 1.47;\ P_{Cl^-} = variable\ (+V_{cmd}) P_{Ca^{2+}} = variable\ (-V_{cmd})$ (negative $V_{cmd}$)

**Cation:** $z_{K^+} = 1;\ c_{K^+cis} = 250\ mM; c_{K^+\,trans} = 25\ mM$

**Anion:** $z_{Cl^-} = -1.0;\ c_{Cl^-cis} = 250mM;\ c_{Cl^-trans} = 25mM$

**Calcium:** $z_{Ca^{2+}} = 2;\ \ c_{Ca^{2+}cis} = 5mM;\ \ \ c_{Ca^{2+}trans} < 10\mu mM$

**Zero-current potential:** $\boldsymbol{Ca^{2+}}: V_{rev} =\approx +47mV\ and\ \boldsymbol{Cl^-}\ V_{rev} =\approx -57mV\ experimental\ values$

Using this values (cis/trans) and the above concentrations in the cis and trans compartment and considering that the assumptions of the GHK-theory are valid under the applied conditions we can use equations (1 to 5 below) to calculate the expected current voltage relation for the above bilayer membrane containing an unknown number of active δ-LIT channels.

Extrapolating the current slope at high positive and negative voltages linearly to zero net current (Figure4b) yields a reversal potential of $V_{rev} = \approx +47mV \ and \ V_{rev} = \approx -57mV$, these values would be compatible with a very high calcium or chloride selectivity of the mature δ-LIT channel depending on the current direction ($P_{Ca^{2+}}/P_{K^+}/P_{Cl^-} \cong 400/1.47/1$) flux of calcium ($-V_{mem}$) or ($P_{Ca^{2+}}/P_{K^+}/P_{Cl^-} \cong 1/1.47/150$) chloride currents ($+V_{mem}$) [31-33]. In summary, the course of the current-voltage relation in Figure 4f can be explained from our calculations if the currents at negative $V_{cmd}$ are carried from cis to trans mainly by $Ca^{2+}$-ions and at positive $V_{cmd}$ predominantly by $Cl^-$ions.

GHK-current equations

1. $I_x(V, P_X, z, c_{cis}, c_{trans}) = P_x z^2 \frac{VF^2}{RT} \cdot \frac{\left(c_{x,cis} - c_{x,trans} exp\left(\frac{-zFV}{RT}\right)\right)}{1 - exp\left(\frac{-zFV}{RT}\right)}$
2. $I_{K^+}(V) = I(V, P_{K^+}, z_{k^+}, c_{K^+cis}, c_{K^+trans}),$
3. $I_{CL^-}(V) = I(V, P_{Cl^-}, z_{Cl^-}, c_{Cl^-cis}, c_{Cl^-trans})$
4. $I_{Ca^{2+}}(V) = I(V, P_{Ca^{2+}}, z_{Ca^{2+}}, c_{Ca^{2+}cis}, c_{Ca^{2+}trans})$
5. $\sum I(V) = I_{K^+}(V) + I_{Cl^-}(V) + +I_{Ca^{2+}}(V)$

Using equation 1-5 we calculated for the ionic conditions given above the corresponding current-voltage relations using a Mathcad (PTC-Software) based routine[34] .
